# Increased and ectopic expression of *Triticum polonicum VRT-A2* underlies elongated glumes and grains in hexaploid wheat in a dosage-dependent manner

**DOI:** 10.1101/2020.11.09.375154

**Authors:** Nikolai M. Adamski, James Simmonds, Jemima F. Brinton, Anna E. Backhaus, Yi Chen, Mark Smedley, Sadiye Hayta, Tobin Florio, Pamela Crane, Peter Scott, Alice Pieri, Olyvia Hall, J. Elaine Barclay, Myles Clayton, John H. Doonan, Candida Nibau, Cristobal Uauy

**Author notes:** The author responsible for distribution of materials integral to the findings presented in this article is: Cristobal Uauy.

## Abstract

Flower development is a major determinant of yield in crops. In wheat, natural variation for the size of spikelet and floral organs is particularly evident in *Triticum polonicum*, a tetraploid subspecies of wheat with long glumes, lemmas, and grains. Using map-based cloning, we identified *VRT2*, a MADS-box transcription factor belonging to the *SVP* family, as the gene underlying the *T. polonicum* long-glume (*P1)* locus. The causal *P1* mutation is a sequence re-arrangement in intron-1 that results in both increased and ectopic expression of the *T. polonicum VRT-A2* allele. Based on allelic variation studies, we propose that the intron-1 mutation in *VRT-A2* is the unique *T. polonicum* species defining polymorphism, which was later introduced into hexaploid wheat via natural hybridizations. Near-isogenic lines differing for the *P1* locus revealed a gradient effect of *P1* across florets. Transgenic lines of hexaploid wheat carrying the *T. polonicum VRT-A2* allele show that expression levels of *VRT-A2* are highly correlated with spike, glume, grain, and floral organ length. These results highlight how changes in expression profiles, through variation in *cis*-regulation, can impact on agronomic traits in a dosage-dependent manner in polyploid crops.

**One-sentence summary:** An intron-1 rearrangement in the MADS-box transcription factor *VRT-A2* leads to its misexpression and defines the long-glume phenotype of Polish wheat (*T. polonicum*).

## Introduction

The genus *Triticum* contains multiple wheat subspecies exhibiting traits of agronomic interest, making them valuable genetic resources for breeding. Among these, *Triticum turgidum* ssp. *polonicum* (Polish wheat), a tetraploid (AABB) spring wheat, is characterized by elongated glumes and grains, the latter of which is an important component of crop yield. Glumes are sterile bract-like organs that subtend spikelets, which are lateral branches that contain several grain-producing florets. Each floret is composed of two leaf-like sheathing structures, the lemma and the palea, as well as two lodicules, three stamens and a pistil (Figure 1A).

**Figure 1.**
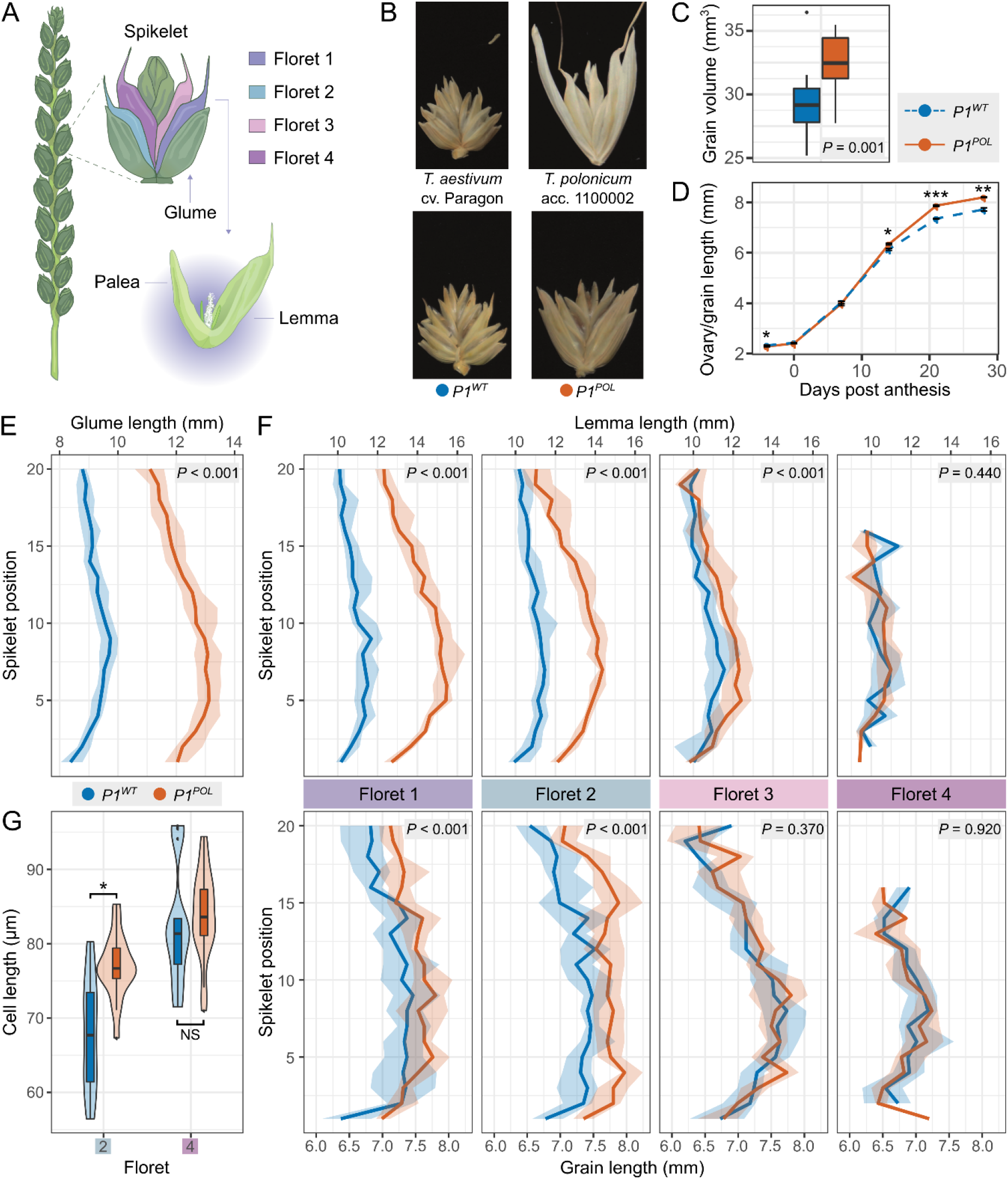
Phenotypic effects of P1 in near-isogenic lines (NILs) (**A**) Drawing of a wheat spike, with a close-up of an individual spikelet. The first four florets on the spikelet are colour coded. A close-up of an open floret depicts its two enveloping sheathing structures (lemma and palea). (**B**) Spikelets of the parental hexaploid bread wheat cultivar ‘Paragon’ and tetraploid *T. polonicum* accession 11000002, and the *P1*^*WT*^ and *P1*^*POL*^ NILs. (**C**) Grain volume was measured using a CT-scanner to image field-grown spikes of the two NILs (n=15). (**D**) Timecourse tracking ovary/grain length development in field-grown *P1*^*WT*^ and *P1*^*POL*^ NILs (n=50). (**E**) Glume length along spikes of *P1*^*WT*^ and *P1*^*POL*^ NILs; positions are numbered from basal (position 0) to apical (position 20) spikelets (n=15 spikes). (**F**) Lemma and grain length at each floret position along *P1*^*WT*^ and *P1*^*POL*^ NILs spikes. Spikelet positions as in E (n= 15 spikes). In (E) and (F) bold line represents the median value, ribbon represents the interquartile range. (**G**) Pericarp cell length from middle sections of grains from floret 2 and floret 4 for the *P1*^*WT*^ and *P1*^*POL*^ NILs (n= 18 grains). In (C) and (G), the box represents the middle 50% of data with the borders of the box representing the 25^th^ and 75^th^ percentile. The horizonal line in the middle of the box represents the median. Whiskers represent the minimum and maximum values, unless a point exceeds 1.5 times the inter-quartile range in which case the whisker represents this value and values beyond this are plotted as single points (outliers). See Supplementary Table S6 for additional measurements. Error bars represent mean ± SEM. *, *P* < 0.05; **, *P* < 0.01; ***, *P* < 0.001.

It was established over 100 years ago that glume length in *T. polonicum* is controlled by a single locus (Engledow, 1920; Biffen, 2009). The *P* or *P1* locus (from <u>P</u>olish wheat) was mapped to chromosome 7A (Matsumura, 1950) and subsequent studies refined the map location to the short arm of chromosome 7A (Watanabe *et al*., 1996; Kosuge *et al*., 2010; Okamoto and Takumi, 2013). While *T. polonicum* as a subspecies is defined by its highly elongated glumes, Biffen (2009), Engledow (1920), and Okamoto and Takumi (2013) also observed that the long-glume trait was completely linked with elongated grains, suggesting multiple pleiotropic effects of the *P1* locus. Okamoto and Takumi (2013) further showed that the *T. polonicum P1* allele was also linked to an increase in spike length and a reduction in the number of spikelets per spike. These studies all determined a semi-dominant effect of *P1*, with heterozygous lines being intermediate to the parents for both glume and grain length.

In addition to tetraploid *T. polonicum*, there are a number of hexaploid bread wheat accessions with elongated glumes. These include the Chinese landrace *T. petropavlovskyi* (also called ‘Daosuimai’ or rice-head wheat) as well as members of the Portuguese landrace group ‘Arrancada’. It is hypothesized that the long-glume phenotype of these hexaploid wheat accessions is the result of natural hybridisation between *T. polonicum* and local landraces (Chen *et al*., 1985; Chen *et al*., 1988; Watanabe and Imamura, 2002; Akond and Watanabe, 2005; Akond *et al*., 2008). Indeed, for both *T. petropavlovskyi* and ‘Arrancada’ the causal genetic locus for long glumes was mapped to chromosome 7A, supporting the hypothesis of a shared origin with *T. polonicum* (Watanabe and Imamura, 2002; Watanabe *et al*., 2004).

The spatial and temporal expression of MADS-box transcription factors determine floral organ identity and developmental phase transitions in plants. The *Tunicate1* mutant of maize (*Zea mays*), known as pod corn, exhibits highly elongated leaf-like glumes that cover the kernels (Mangelsdorf and Galinat, 1964; Langdale *et al*., 1994). Genetic studies identified the causal gene as *Zea mays MADS19* (*ZMM19*), a member of the *short vegetative phase* (*SVP*) gene family of MADS-box transcription factors (Han *et al*., 2012; Wingen *et al*., 2012). A rearrangement in the promoter region of *ZMM19* causes its ectopic expression, which leads to the dosage-dependent phenotype (Han *et al*., 2012; Wingen *et al*., 2012). Ectopic expression of *ZMM19* in *Arabidopsis thaliana* leads to enlarged sepals, suggesting a conserved mechanism (Wingen *et al*., 2012).

Spikelet morphology and organ size are tightly correlated with final grain weight in wheat (Millet, 1986). Despite their importance, we have relatively little understanding of the genes controlling spikelet and floral organ size in wheat. Here, we characterised the *P1* locus of *T. polonicum*, which has pleiotropic effects on glume, floral organ, and grain size. We show that the *P1* long-glume phenotype is due to the ectopic expression of *VRT-A2*, a *SVP* MADS-box transcription factor. The higher and ectopic expression of *VRT-A2* is due to a sequence rearrangement in the first intron, which defines *T. polonicum* as a subspecies. Expression levels of *VRT-A2* affect glume, grain, and floral organ length in a dosage-dependent manner.

## Results

### The long-glume T. polonicum P1 allele enhances grain weight through longer grains

To evaluate the performance of the *T. polonicum P1* allele we developed BC_4_ and BC_6_ near isogenic lines (NILs) by crossing *T. polonicum* accession 1100002 to the hexaploid spring wheat cultivar Paragon (Table 1, Figure 1B). We verified the isogenic status of these lines using the Breeders’ 35K Axiom Array (Supplementary Figure S1; (Allen *et al*., 2017)) and assessed the *T. polonicum* (*P1*^*POL*^) and wildtype (*P1*^*WT*^) NILs in the field over multiple years and environments. The *P1*^*POL*^ NILs had longer glumes and lemmas than wildtype Paragon NILs (Figure 1B) and were on average 6 cm taller due to an increase in peduncle (final internode) and spike lengths (1.6 cm; *P* <0.01; Table 1, Supplementary Table S1-S3, Supplementary Figure S2). The increase in spike length, alongside a minor increase in spikelet number (0.4 spikelets per spike; *P* = 0.06), led to a 11.3% decrease in spikelet density (spikelets per cm) in *P1*^*POL*^ with respect to *P1*^*WT*^ NILs (significant in three of four environments; Supplementary Table S1). The *P1*^*POL*^ NILs also flowered on average 0.8 days later (*P* <0.001) than the wildtype NILs. We observed consistent positive effects on thousand grain weight in *P1*^*POL*^ (TGW; 5.5%; *P* <0.001), which were driven by significant increases in grain length (5.0%; *P* <0.001), but not grain width (Table 1, Supplementary Table S1, S4). The increase in grain length resulted in an increased grain volume (10.7%; *P* = 0.001) as determined by CT scans of a single year of field samples (Figure 1C, Supplementary Table S4). The increased TGW in *P1*^*POL*^ NILs also translated into a significant increase in hectolitre weight (HLW) in 4 out of 5 environments (2.3%; *P* <0.01). Final yield, however, was not significantly different between NILs despite the increase in TGW (Table 1, Supplementary Table S1).

**Table 1.**
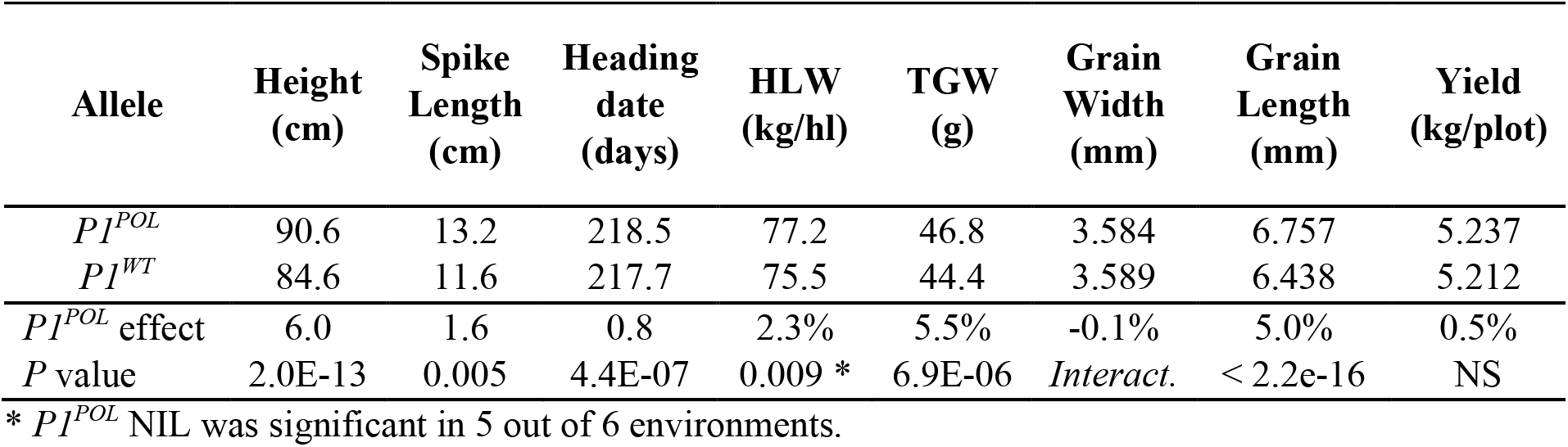
**Phenotypic effects of the *P1* allele in Paragon NILs**. *P1*^*POL*^ effect is the percentage difference (except height and spike length in cm; heading date in days) between the *P1*^*WT*^ and the *P1*^*POL*^ NILs. The *P* value of the ANOVA main effect is presented, apart from grain width, which had a significant interaction across environments (simple effects and detailed breakdown in Supplementary Table S1). Values represent means of six field experiments (except spike length n=4).

### The P1 allele from T. polonicum enhances grain size in florets 1 and 2 through an increase in cell length

We conducted more in-depth phenotyping to identify the first timepoint during grain development in which differences in grain length are established between *P1* NILs. We dissected and measured field-grown grain samples from florets 1 and 2 of five central spikelets from *P1*^*WT*^ and *P1*^*POL*^ NILs at six timepoints during grain development (Figure 1D; second year data in Supplementary Figure S3). We did not detect consistent differences in ovary length before and at anthesis nor in grain length 7 days post anthesis (dpa). However, at 14 <u>d</u>pa, grains from *P1*^*POL*^ NILs were 3.4 % longer than grains from *P1*^*WT*^ NILs (*P* <0.05; Supplementary Table S5). The increased grain length in *P1*^*POL*^ NILs was maintained at 21 and 28 dpa (7.0 and 6.2% longer grains than *P1*^*WT*^, respectively; *P* <0.002; Supplementary Table S5). These results, consistent in two independent field seasons (Supplementary Figure S3), suggest that the difference in grain length between *P1* NILs is established during mid-grain filling.

We next measured the size (length, width, and area) of glumes, floral organs (lemma and palea), and grains across spikes and spikelets of *P1*^*POL*^ and *P1*^*WT*^ NILs using the same field-grown samples as above. We focus on organ length, given the effects observed in the field (Table 1, Fig. 1D), and all data, including width and area measurements, are presented in Supplementary Tables S2-S3 and Supplementary Figures S4-S5. The *T. polonicum P1* allele significantly increased glume length (37%; *P* <0.001) with respect to the wildtype NILs; this effect was consistent and independent of spikelet position across the spike (Figure 1E). However, we detected a significant gradient in the effect of the *T. polonicum P1* allele across the florets within each spikelet: the largest and most significant effects on lemma length were observed in florets 1 and 2 (28.6% and 19.8%, respectively; *P* <0.001), whereas the effect was reduced in floret 3 (+5.8%; *P* <0.001) and was non-significant in floret 4 (Figure 1F, Supplementary Table S3). This gradient in lemma length within spikelets was maintained across all positions along the spike. A very similar gradient within spikelets was also identified for grain length, with *P1*^*POL*^ NILs having significantly longer grains than *P1*^*WT*^ NILs in florets 1 and 2 (+4.1% and +6.7%, respectively; *P* <0.001), and non-significant differences in grain length in florets 3 (+0.4%; *P*=0.37) and 4 (+0.1%; *P*=0.92; Figure 1F). Minor effects of the *T. polonicum P1* allele on palea length also followed this spikelet gradient (+2.5% in floret 1 to -3.6% in floret 4; Supplementary Table S3, Supplementary Figure S5). These results suggest that the increases in glume, lemma, and grain length conferred by the *T. polonicum P1* allele are consistent along the spike, but that the positive effects on lemma and grain length follow a basipetal gradient from basal to apical florets within individual spikelets.

To further investigate the differences in grain length between *P1* NILs, we used scanning electron microscopy to image and measure pericarp cell size of *P1*^*WT*^ and *P1*^*POL*^ grains. We selected grains from florets 2 and 4 of central spikelets and imaged the base, centre, and distal end of each grain (Supplementary Figure S6A). We found a significant 12.9 % increase (*P* <0.05) in pericarp cell length in floret 2 grains of *P1*^*POL*^ NILs relative to *P1*^*WT*^ NILs (Figure 1G). This difference in cell length was present only in the central portion of the grain, while cell size was similar between NILs at the base and distal end of the grain (Supplementary Table S6, Supplementary Figure S6). For floret 4, there were no differences in pericarp cell size between the NILs at each of the three positions examined across the grain (Figure 1G, Supplementary Table S6, Supplementary Figure S6). Given the maternal origin of the pericarp, these results are consistent with the *P1*^*POL*^ grain length effect being maternally inherited as first proposed by Engledow (1920). Taken together, these results suggest that the *T. polonicum P1* allele enhances grain size in basal florets through an increase in cell length in the centre of the grain, whereas grains of floret 3 and 4 are indistinguishable from the wildtype NILs, both macro- and microscopically.

### P1 maps to a 50 kb interval on chromosome 7A containing a single candidate gene

To map the *P1* locus, we used BC_4_ and BC_6_ recombinant lines derived from the NILs described above. We initially phenotyped 17 BC_4_F_3_ homozygous recombinant lines between markers *S1* and *S9* for glume length and mapped the *P1* locus between markers *S2* (125,260,256 bp) and *S7* (150,240,183 bp) (Figure 2A, Supplementary Table S7). Heterozygous individuals across the interval had glumes of intermediate length between the homozygous parental lines, consistent with a semi-dominant mode of action of *P1* (Engledow, 1920; Biffen, 2009; Okamoto and Takumi, 2013). To further define the *P1* interval, we identified an additional 64 homozygous BC_6_F_2_ recombinants between markers *S2* and *S10*, which were genotyped with a further 21 markers (Figure 2B, Supplementary Table S8). The long-glume phenotype, alongside plant height, spike length, grain length, and thousand grain weight, mapped between markers *S15* and *S19*, spanning a 50,338 bp interval (Figure 2C, Supplementary Table S8-S9). The complete linkage of the 50.3 kbp region with these multiple phenotypes suggests that they are all pleiotropic effects of the *P1* locus.

**Figure 2.**
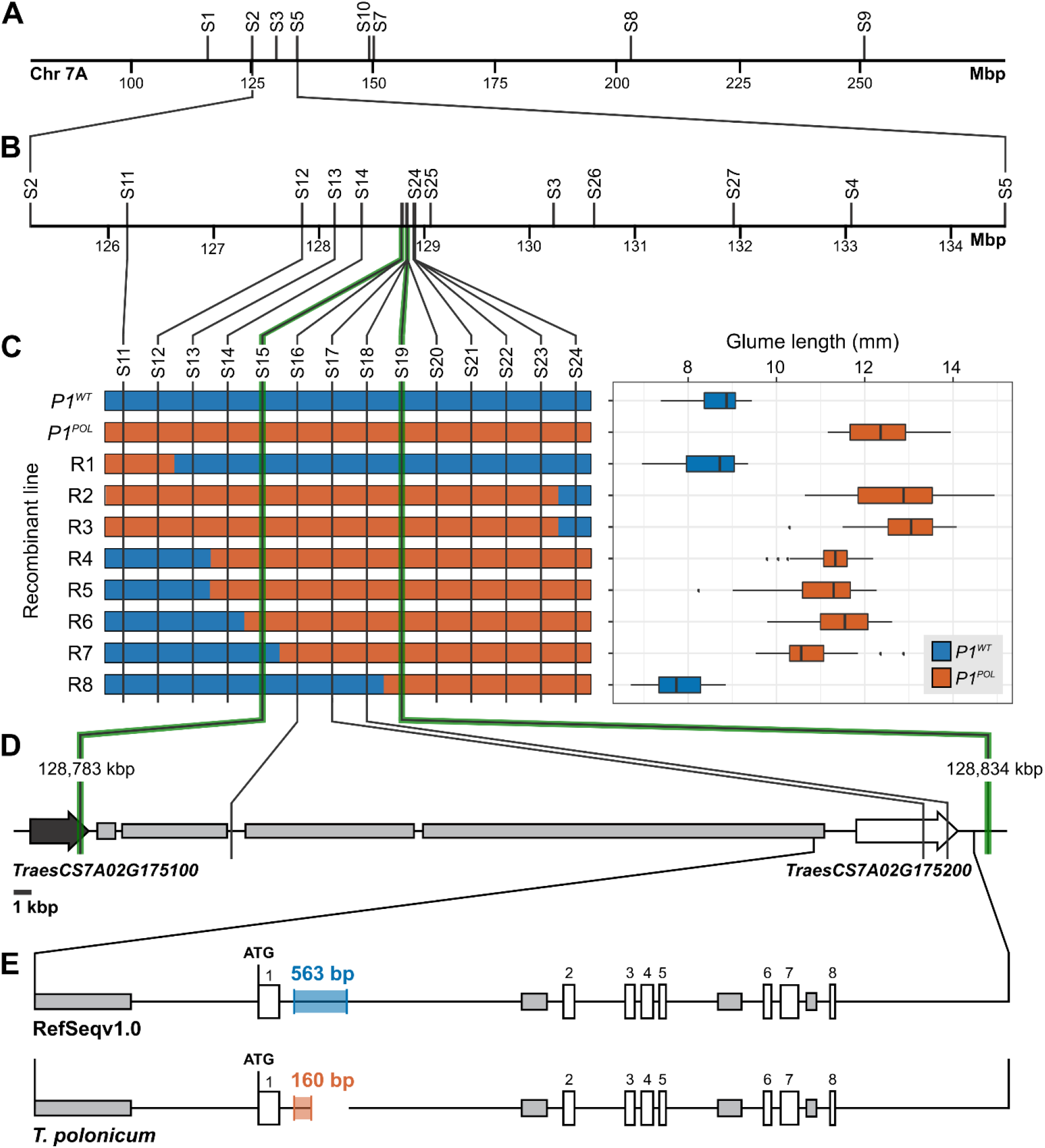
Map-based cloning of the P1 locus reveals VRT-A2 as the single candidate gene. (**A**) Initial mapping in 17 BC_4_F_2_ recombinants mapped *P1* between markers *S2* and *S7* (∼25 Mbp interval). (**B**) Subsequently, *P1* was mapped between markers *S2* and *S10* using an additional 64 BC_6_F_2_ recombinant lines. (**C**) Graphical genotype of eight critical recombinants between markers *S12* and *S24* (∼1 Mbp interval; marker distance not drawn to scale). Based on the phenotypic evaluation of glume length, we mapped *P1* to a 50.3 kbp interval between markers *S15* and *S19* (n=5 plants per genotype). The box represents the middle 50% of data with the borders of the box representing the 25^th^ and 75^th^ percentile. The horizonal line in the middle of the box represents the median. Whiskers represent the minimum and maximum values, unless a point exceeds 1.5 times the inter-quartile range in which case the whisker represents this value and values beyond this are plotted as single points (outliers). (**D**) The 50.3 kbp interval encompasses the last exon of *Traes7A02G175100* (black arrow), multiple repetitive elements (grey rectangles) and *Traes7A02G175200* (red arrow). (**E**) We identified a single polymorphism between Chinese Spring (RefSeqv1.0) and *T. polonicum* in a ∼10 kbp interval surrounding *Traes7A02G175200*. A 563-bp sequence in RefSeqv1.0 was substituted by a 160-bp sequence in *T. polonicum*.

We identified two gene models based on the RefSeqv1.1 annotation within the *P1* interval: *TraesCS7A02G175100* and *TraesCS7A02G175200*. The flanking marker *S15* resided within the last intron of *TraesCS7A02G175100* and no additional SNPs were detected in the last exon of this gene between *P1*^*WT*^ and *P1*^*POL*^ NILs. Manual annotation of the 50.3 kbp *P1* interval in the RefSeqv1.0 assembly (and an additional 14 hexaploid and tetraploid cultivars) identified 38,261 bp as repetitive sequences, and no additional gene apart from *TraesCS7A02G175200* (Figure 2D, Supplementary Figure S7A). This suggested *TraesCS7A02G175200* as the sole candidate gene for *P1*.

*TraesCS7A02G175200* encodes a member of the MADS-box gene family previously named *VEGETATIVE TO REPRODUCTIVE TRANSITION 2 (VRT2)* in wheat (Kane *et al*., 2005). *VRT2*, as well as its homolog *TaSVP1*, are the wheat orthologs of *AtSVP* in *Arabidopsis thaliana* (Supplementary Figure S8; (Schilling *et al*., 2020)). Using publicly available RNA-Seq data, we verified the exonintron structure of *TraesCS7A02G175200*.*1* (Supplementary Figure S7B). We sequenced the gene in the *P1*^*POL*^ NIL from 2299 bp upstream of the ATG to marker *S19* (1857 bp downstream of the termination codon; 9747 bp total including all exons and introns). Compared to the RefSeqv1.0 assembly, we only found a single polymorphism located within intron-1; a 563-bp sequence in the RefSeqv1.0 assembly that was substituted for a 160-bp sequence in *P1*^*POL*^ (Figure 2E, Supplementary Table S10). An analysis of the 160-bp sequence suggests that it consists of reoccurring units of DNA which are homologous to sequences found in the intron 1 regions flanking either side of the 160-bp substitution (Supplementary Figure S9). These reoccurring sequence units account for 145 of the 160 nucleotides, indicating that the sequence is the result of a local rearrangement rather than insertion of foreign DNA.

### The 160-bp intron-1 sequence rearrangement in VRT-A2 is completely linked with the long-glume phenotype in tetraploid and hexaploid wheat accessions

We determined the allelic status of the *VRT-A2* intron-1 sequence rearrangement in a wheat diversity panel. We first screened 367 accessions with wildtype glume length including tetraploid emmer wheat *T. dicoccoides* (n=70), tetraploid durum wheat *T. durum* (n=21), hexaploid wheat landraces from the Watkins collection (n=103), hexaploid UK cultivars (n=98), hexaploid European germplasm from the Gediflux collection (n=60), and 15 sequenced wheat cultivars. We found that all 367 accessions carried the 563-bp sequence in intron 1 and none had the 160-bp sequence rearrangement found in the *P1*^*POL*^ NIL (Supplementary Table S11). We thus termed the 563-bp sequence as the wildtype *VRT-A2a* allele, and the 160-bp sequence rearrangement found in *T. polonicum* as the *VRT-A2b* allele, consistent with wheat gene nomenclature guidelines. We next screened 36 accessions of tetraploid *T. polonicum* (all with long glumes) from 17 different countries to determine their *VRT-A2* allele. All 36 *T. polonicum* accessions carried the exact 160-bp rearrangement in intron 1 as the *VRT-A2b* allele (Figure 3A, B, Supplementary Table S12). These results suggest that the *VRT-A2b* allele with its 160-bp sequence rearrangement in intron 1 is unique to *T. polonicum*.

**Figure 3.**
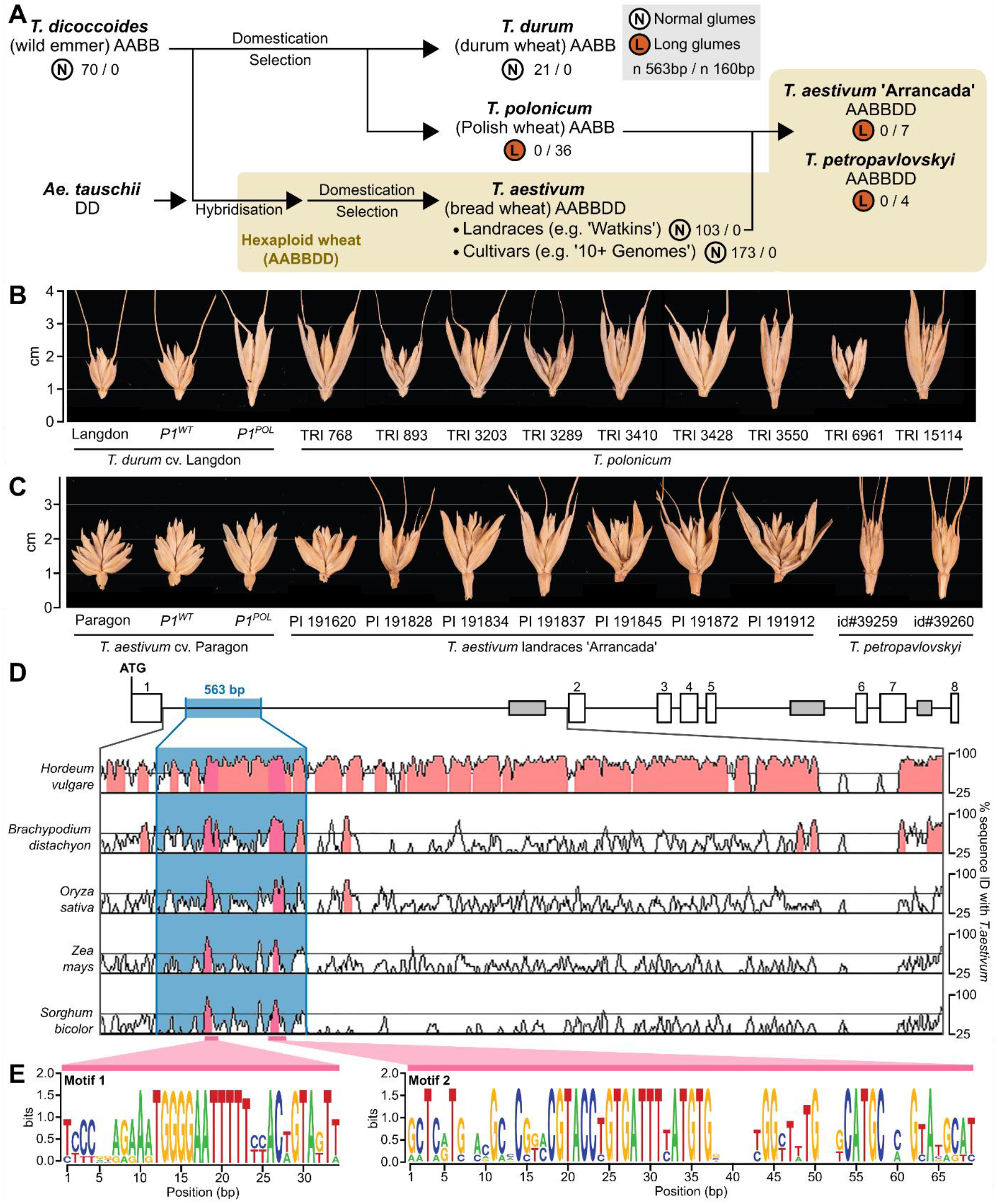
Natural variation of P1 indicates a single mutation event that led to the loss of evolutionary conserved motifs. (**A**) Simplified diagram depicting the evolution and domestication of tetraploid and hexaploid wheat (tan shaded area). The glume phenotype for each species and set of accessions is indicated by the letter N (normal) or L (long) enclosed in a circle. Beside this classification, the number of accessions that carry the wildtype 563-bp (*VRT-A2a* allele) or the 160-bp (*VRT-A2b* allele) intron 1 sequence is shown. *T. polonicum* hybridised with hexaploid landraces in China and Portugal, giving rise to *T. petropavlovskyi* and the ‘Arrancada’ landrace group, respectively, both of which exhibit long glumes and carry the *VRT-A2b* allele. (**B**) Spikelets of tetraploid wheat including *P1* NILs in the tetraploid cultivar ‘Langdon’ and nine accession of *T. polonicum*. (**C**) Comparison of spikelets of hexaploid wheat including *P1* NILs in the hexaploid cultivar ‘Paragon’, seven accession from the ‘Arrancada’ landrace group and two accession of *T. petropavlovskyi*. (**D**) Phylogenetic shadowing using mVISTA of *VRT-A2* intron 1 with pairwise alignments of *T. aestivum* with barley (*Hordeum vulgare*), *Brachypodium distachyon*, rice (*Oryza sativa*), maize (*Zea mays*), and sorghum (*Sorghum bicolor*). The Y-axis represents percentage sequence similarity. Two conserved peaks (dark pink) were identified within the 563-bp sequence (blue box) that is absent in *T. polonicum*. (**E**) Sequence of the two conserved motifs that maintain an >80% similarity over a 20 bp sliding window across the species described in D.

We then examined accessions from two types of hexaploid wheat with long-glumes that have been postulated to be the product of independent hybridisation between *T. polonicum* and hexaploid landraces in China (*T. petropavlovskyi*) and Portugal (‘Arrancada’ group; Figure 3A, C; (Chen *et al*., 1985; Chen *et al*., 1988; Watanabe and Imamura, 2002; Akond and Watanabe, 2005; Akond *et al*., 2008)). All 11 accessions of *T. petropavlovskyi* (n=4) and the ‘Arrancada’ landraces (n=7) carried the *VRT-A2b* allele found in *T. polonicum* (Supplementary Table S12). We fully sequenced the allele (5591 bp) from six *T. polonicum*, two *T. petropavlovskyi*, and four ‘Arrancada’ accessions (Supplementary Table S12) and obtained 100% identical sequences from these twelve long-glumed accessions. Using the markers developed for mapping *P1*, we found that our two *T. petropavlovskyi* accessions shared a common haplotype, whereas the seven ‘Arrancada’ accessions also shared a common, albeit distinct, haplotype from that in *T. petropavlovskyi* (Supplementary Table S13). Conversely, in accessions with normal-sized glumes, we identified multiple haplotypes within *VRT-A2* (all with the 563-bp intron-1 sequence) and also across the wider physical interval (Supplementary Tables S10, S13). These results, alongside the absence of the 160-bp rearrangement in wild emmer and hexaploid landraces, provide evidence that the 563-bp intron-1 sequence in *VRT-A2a* is ancestral.

### The 563-bp intron-1 sequence of VRT-A2a is highly conserved across Poaceae

We compared the entire intron-1 sequence of *VRT-A2a* with orthologous Poaceae sequences from barley (*Hordeum vulgare*), *Brachypodium distachyon*, rice (*Oryza sativa*), maize (Zea mays), and sorghum (*Sorghum bicolor*). Phylogenetic shadowing using mVISTA (Mayor *et al*., 2000; Frazer *et al*., 2004) revealed two highly conserved regions across Poaceae (>85% sequence id, minimum 20 bp), both of which are missing from the 160-bp rearrangement present in the *VRT-A2b* allele (Figure 3D). We further examined these two regions (see Methods) and identified broadly conserved sequences of 34 and 69 bp in length, hereafter referred to as ‘Motif 1’ and ‘Motif 2’ respectively (Figure 3E). Within them, both motifs contain highly conserved sequences of 16 and 20 bp, respectively (Supplementary Data Set S1). We searched for putative transcription factor binding sites within Motifs 1 and 2 using three online databases (PlantPan3.0, PlantRegMap, and MEME, see Methods). We found two significant hits (both *P* <0.001 and *q* <0.001) for Motif 1 encoding members of the LATERAL ORGAN BOUNDARIES-DOMAIN (LBD) family (Supplementary Data Set S2). For Motif 2, we found 17 significant hits (all *P* <0.001 and *q* <0.05) encoding members of the LBD, Basic Leucine Zipper (bZIP), B3, and GLABROUS1 enhancer-binding protein (GeBP) families (Supplementary Data Set S2). Given the highly conserved nature of motifs within the 563-bp intron-1 sequence across the investigated Poaceae (∼60 million years divergence time; (Charles *et al*., 2009; Reineke *et al*., 2011)) and the identification of putative transcription factor binding sites, we hypothesise that this intron-1 sequence plays a regulatory role in the expression profile of *VRT-A2*.

### VRT-A2 is expressed ectopically and more highly in P1^POL^ relative to P1^WT^ NILs

To assess if the intron-1 sequence rearrangement affected the expression profile of *VRT-A2*, we determined its expression pattern in *P1*^*WT*^ and *P1*^*POL*^ NILs using qRT-PCR. We first examined expression levels in developing meristems of *P1*^*WT*^ NILS. Consistent with previous studies (Kane *et al*., 2005; Kane *et al*., 2007; Trevaskis *et al*., 2007), we found a progressive decrease in *VRT-A2* expression from vegetative meristem (W1) to terminal spikelet stage (W4; Figure 4A). We next examined expression in *P1*^*POL*^ NILs, which showed a five to 16-fold higher expression level of *VRT-A2* relative to the *P1*^*WT*^ NILs at all five timepoints investigated (*P* <0.05; Figure 4A). An increased expression level of *VRT-A2* in *P1*^*POL*^ relative to *P1*^*WT*^ was also observed in leaves at the same developmental stages (six to 38-fold higher, *P* <0.05; Supplementary Table S14). Next, we examined *VRT-A2* expression in developing glumes, lemmas, anthers, flag leaves, and grains at multiple developmental timepoints. In the wildtype NIL, expression was restricted to the flag leaves and anthers at anthesis (Figure 4B, Supplementary Table S14). We did not detect *VRT-A2* expression in glumes, lemmas, nor grains of *P1*^*WT*^ NILs at any timepoint (Figure 4B-D), consistent with publicly available RNA-Seq data of wildtype *VRT-A2* genotypes (Borrill *et al*., 2016; Ramirez-Gonzalez *et al*., 2018). By contrast, *VRT-A2* expression was detected in all tissues and at all timepoints in the *P1*^*POL*^ NIL, including glumes, lemmas, and grains (Figure 4B-D).

**Figure 4.**
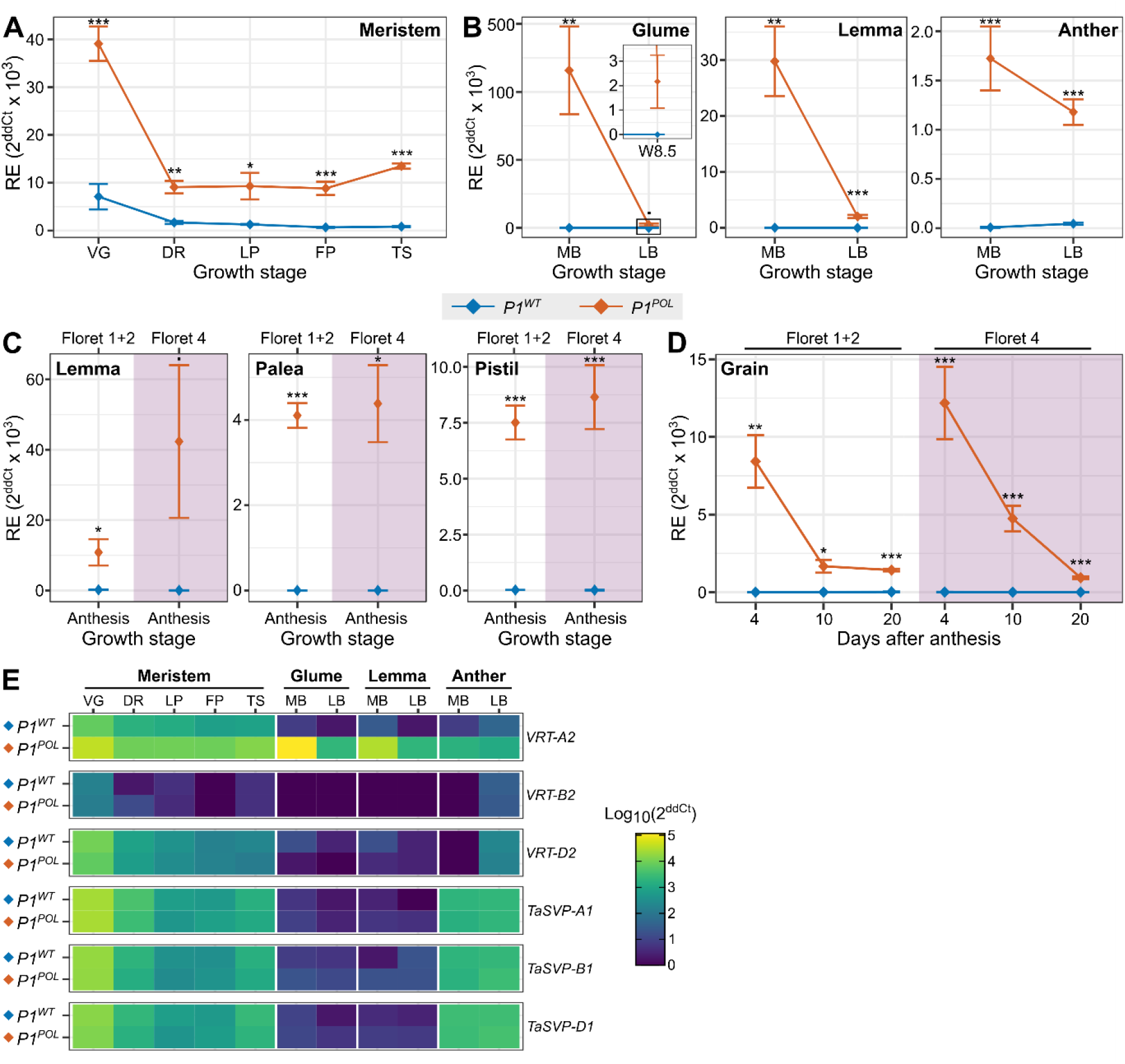
VRT-A2 is more highly and ectopically expressed in P1^POL^ relative to P1^WT^ NILs. (**A**) Relative expression of *VRT-A2* in developing meristems of *P1*^*WT*^ (blue) and *P1*^*POL*^ (orange) NILs. Developmental stages based on Waddington scale (Waddington *et al*., 1983); VG vegetative meristem (W1); DR, late double ridge stage (W2.5); LP, lemma primordium stage (W3.25); FP, floret primordium stage (W3.5); TS, early terminal spikelet stage (W4). (**B**) Relative expression of *VRT-A2* in glume, lemma (floret 1+2), and anther (floret 1+2) at mid-boot (W7.5) and late-boot (W8.5) stages. (**C**) Relative expression of *VRT-A2* in lemma, palea, and pistil just before anthesis (W9.5) in florets 1+2 (white background) and floret 4 (pink background). Note that for Panel C and D the growth stage is based on florets 1+2; floret 4 tissues will be at a slightly less mature developmental stage. (**D**) Relative expression of *VRT-A2* in grains from florets 1+2 as well as floret 4 at 4, 10, and 20 days post anthesis. (**E**) Heatmap showing log_10_ scaled expression in *P1*^*WT*^ and *P1*^*POL*^ NILs for the three *VRT2* and *TaSVP1* homoeologs in tissues and developmental stages shown in panels A and B (Supplementary Table S14). Error bars are mean ± SEM. *, *P* < 0.05; **, *P* < 0.01; ***, *P* < 0.001.

Given the contrasting effects of the *VRT-A2b* allele on grain and lemma length between floret 1+2 and floret 4 (Figure 1F), we compared the *VRT-A2* ectopic expression in these samples. Across multiple tissues and timepoints (lemma, palea, pistil at anthesis; grains at 4, 10, and 20 days post anthesis) we found similar expression of *VRT-A2* in samples from florets 1+2 compared to floret 4 (Figure 4 C, D). We also investigated expression of the *VRT2* homoeologs (*TraesCS7B02G080300, TraesCS7D02G176700*), and that of the closely related ortholog *TaSVP1* (*TraesCS6A02G313800, TraesCS6B02G343900, TraesCS6D02G293200*) (Supplementary Figure S8; (Schilling *et al*., 2020)). Across the same tissues and developmental timepoints as those described above, we did not detect any differences in expression between *P1*^*WT*^ and *P1*^*POl*^ NILs (Figure 4E). These results show that *VRT-A2* is expressed more highly and ectopically in *P1*^*POL*^ relative to *P1*^*WT*^ NILs across multiple tissues and timepoints. The ectopic expression does not extend to the homoeologs or closely related orthologs and is not restricted to those tissues in which we observed phenotypic differences between *P1*^*WT*^ and *P1*^*POL*^ NILs.

#### Ectopic expression of VRT-A2 leads to phenotypic effects in a dosage dependent manner

We next tested whether the observed changes in *VRT-A2* expression patterns in *P1*^*POL*^ NILs are causal for the *T. polonicum* long-glume phenotype. We transformed the hexaploid accession ‘Fielder’ (normal glume phenotype) using the genomic *T. polonicum VRT-A2b* allele (5591 bp), including 2299 bp upstream of the ATG, all coding and intron sequences, as well as 1000 bp downstream of the termination codon. We obtained 14 independent T_0_ lines, which were classified based on the transgene copy number. No transgene was detected for five lines (zero copy number), three lines carried 1 or 2 copies (low copy number), three lines carried 4 to 5 copies (medium copy number), and three lines carried 9 to 35 copies (high copy number; Figure 5A, Supplementary Tables S15-S17).

**Figure 5.**
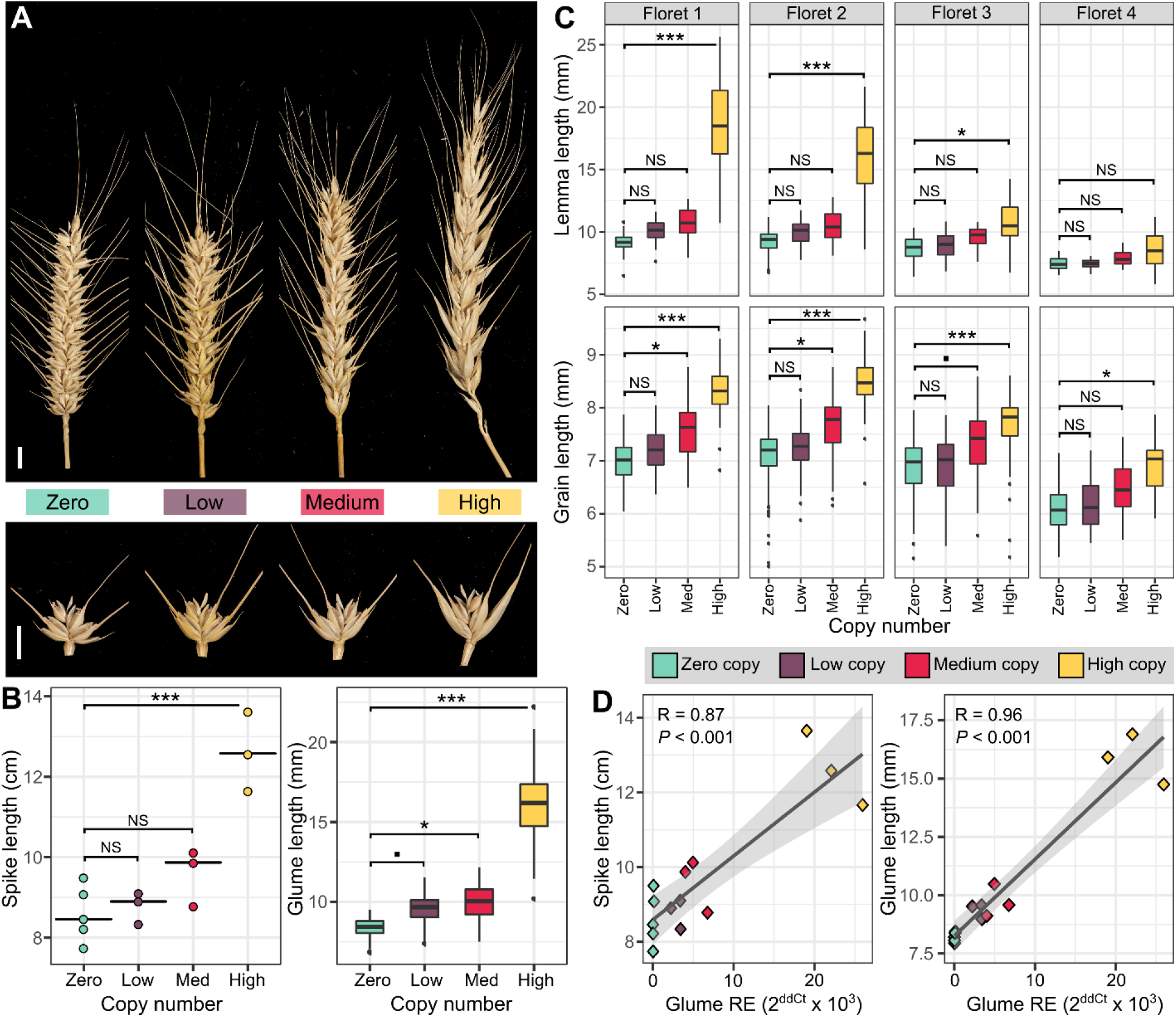
Increased and ectopic VRT-A2 expression elicits phenotypic effects in a dosage dependent manner. (**A**) Comparison of spikes and spikelets of zero, low (1-2), medium (3-4), and high (9-35) copy number lines (left to right). Notably, spike length increases with copy number, as does glume length. Scale bar = 1 cm. (**B**) Dot and box plots depicting the variation of spike (left) and glume (right) length, respectively, from two tillers of zero (cyan), low (purple), medium (red), and high (yellow) copy number lines. Horizontal lines represent the median. (**C**) Box plots depicting lemma and grain length for florets 1, 2, 3, and 4 for zero (cyan), low (purple), medium (red), and high (yellow) copy number lines. (**D**) Pearson correlations between *VRT-A2* relative expression in the glume at 21 dpa and spike length (left), and glume length (right). Relative expression shown as 2^ddCt^ x 10^3^ (Supplementary Table S15). Regression (Dark grey line) and 95% confidence interval (light grey shading) are shown. Data points are coloured according to copy number. Additional correlations in Supplementary Table S17. Box plots in (B) and (C) include all subsamples, whereas statistical analyses were performed with mean values. The box represents the middle 50% of data with the borders of the box representing the 25^th^ and 75^th^ percentile. The horizonal line in the middle of the box represents the median. Whiskers represent the minimum and maximum values, unless a point exceeds 1.5 times the inter-quartile range in which case the whisker represents this value and values beyond this are plotted as single points (outliers). Statistical classifications in (B) and (C) are based on Dunnett tests against the zero copy number lines. ▪, *P* < 0.10; *, *P* < 0.05; **, *P* < 0.01; ***, *P* < 0.001.

We collected tissue from flag leaves, glumes, and grains at 21 dpa from all 14 plants to measure expression levels of *VRT2* homoeologs (Supplementary Table S15). We detected expression of *VRT-A2* in flag leaves of all 14 transformed lines, including the zero copy number lines, similar to that observed in *P1*^*WT*^. In glume and grain tissue, *VRT-A2* expression was extremely low or not detected in the zero copy number lines, consistent with the *P1*^*WT*^ NILs (Figure 4), whereas we detected expression in all lines with at least one copy of the transgene. In all tissues, *VRT-A2* expression scaled with copy number. As seen in the NILs, we did not detect differences in expression of the B- and D-homoeologs among transgenic lines (Supplementary Table S15).

We dissected all spikelets from two spikes of each of the 14 T_0_ lines for morphological characterisation. We compared spike length and glume length, as well as lemma, palea, and grain length from florets 1 to 4 among the four categories of copy number lines (Figure 5A-C, Supplementary Table S16). Overall, we identified significant differences between the zero copy lines (N = 5) and the transgenic lines (low, medium, and high; N =9) for glume, lemma, palea, and grain length (all *P* <0.05; Supplementary Table S16). These differences were largest in glumes (42%) and lemmas (26%), whereas paleae showed the smallest effect (6.6%; Figure 5B, C, Supplementary Table S16).

Having established the overall positive effects of the *T. polonicum VRT-A2* transgene on these traits, we next evaluated the magnitude of the phenotypic effects among the four categories of copy number lines. Due to the relatively small sample size for each category (N=3 to 5 T_0_ plants), we did not detect significant differences between zero copy number lines and low copy number lines for any of the traits, although glume length increased by 14.1% (*P* <0.08). However, we did detect significant effects for glume length (19.0%), palea length (5.9%) and grain length (9.6%) in medium copy number lines (all *P* < 0.05). These effects increased in magnitude and significance in high copy number lines, which showed a highly significant increase of 93% in glume length, 58% in lemma length, 13% in palea length, and 14% in grain length with respect to the zero copy number lines (all *P* <0.002; Supplementary Table S16). While spike length was increased in both low and medium copy number lines (3% and 14%, respectively), it was only significantly increased in high copy number lines with a 51% increase compared to zero copy number lines (*P* < 0.01). This significant increase in spike length led to a significant 47% reduction in spikelet density in the high copy number lines, consistent with the effects observed in the *P1*^*POL*^ NILs (Supplementary Table S1). We saw a gradient in the phenotypic effects from floret 1 to floret 4, consistent again with what we observed in the NILs. This basipetal gradient was most obvious in the medium and high copy number lines. For example, floret 1 lemma and grain length increased by 105% and 19%, respectively, in high copy number lines, whereas in floret 4 both traits ‘only’ increased by 11% (Supplementary Table S16).

We further analysed the phenotypic data to relate it to the expression of *VRT-A2* in the different transgenic lines. *VRT-A2* expression correlated highly and significantly with several phenotypic traits, independent of the tissue examined for expression. For example *VRT-A2* expression levels in the glume correlated strongly with spike length (R^2^ = 0.76; *P* <0.0001) and glume length (R^2^ = 0.92; *P* <0.0001) (Figure 5D), as well as with lemma, palea, and grain length in floret 1 (all R^2^ >0.81; all *P* <0.0001; Supplementary Table S17). Similar to the gradient in phenotypic effects across florets detailed above, these correlations were strongest and most significant in floret 1 and floret 2 (R^2^ >0.64), remained significant for floret 3 phenotypes (R^2^ ≥0.41), and were less significant for floret 4 phenotypes (Supplementary Table S17). The multiple phenotypes of the medium and high copy transgenic lines recreate, in a dosage dependent manner, the effects seen in the *P1*^*POL*^ NIL, providing further evidence that *VRT-A2* is the causal gene underlying the *T. polonicum P1* locus.

## Discussion

### An intron-1 sequence rearrangement in VRT-A2 is the T. polonicum species defining polymorphism

The first formal classification of wheat was compiled by Linnaeus in 1753 and was based on discernible characteristics such as phenology, spike architecture, and glume morphology. With its characteristic long glumes *T. polonicum* is a standout *Triticum* species. Despite this, the first mention of *T. polonicum* only dates back to 1687 (Percival, 1921). *T. polonicum* was instrumental in the study of early geneticists who showed that measurable quantitative traits, such as glume length, were also inherited according to the same laws of qualitative traits postulated by Mendel (Engledow, 1920; Biffen, 2009). However, despite first being described genetically over a century ago, the gene underlying the *P1* locus remained unknown.

Here, we show that the gene underlying the long-glume *P1* locus of *T. polonicum* is *VRT-A2*, a member of the *SVP* family of MADS-box transcription factors. We mapped multiple phenotypes associated with *P1*, including glume length, grain length, spike length, grain weight, and plant height to the same physical interval that included *VRT-A2* as the single candidate gene. For grains, we further showed that the increase in length is likely a result of increased cell length. We identified a sequence rearrangement in the first intron of *VRT-A2* in which a 563-bp wildtype sequence (*VRT-A2a*) was replaced by a 160-bp fragment in *T. polonicum* (*VRT-A2b*). The 160-bp sequence is mostly composed of imperfect tandem copies of two sequence units that flank either side of the rearranged intron-1 region (Supplementary Figure S9). The 160-bp sequence has no match to other plant sequences nor to repetitive elements consistent with its local origin rather than an insertion of foreign sequence. We hypothesise that the rearrangement was caused by erroneous DNA repair following a double strand break via the alternative non-homologous end-joining (aNHEJ) pathway (Deriano and Roth, 2013; Rodgers and McVey, 2016). aNHEJ has been shown to result in complex insertion/deletion events with insertions being identical or near-identical matches of flanking sequences (Kent *et al*., 2016; van Kregten *et al*., 2016). These templated insertion are associated with many disease-causing genomic rearrangements in humans (Schimmel *et al*., 2019).

Using diverse germplasm, we showed that the *VRT-A2b* allele is only present in tetraploid *T. polonicum* accessions, or in hexaploid wheat germplasm with long glumes. Our results provide strong evidence that these hexaploid wheat accessions with long-glumes, namely *T. petropavlovskyi* and the ‘Arrancada’ accessions, are the outcome of introgressions between *T. polonicum* and hexaploid wheat (Chen *et al*., 1985; Chen *et al*., 1988; Watanabe and Imamura, 2002; Akond and Watanabe, 2005; Akond *et al*., 2008). The lack of sequence variation among lines with the *VRT-A2b* allele, coupled with its absence among ancestral wheat types (e.g. wild emmer, Watkins landraces), suggests a single and recent origin for the *VRT-A2b* allele. We propose that a single mutation event in the ancestral *VRT-A2a* intron-1 sequence gave rise to the 160-bp sequence rearrangement within the domesticated tetraploid gene pool. This mutation was later introduced into hexaploid wheat via natural hybridizations resulting in hexaploid accessions with long-glumes. These results, together with the complete linkage of this *VRT-A2b* allele with the long-glume phenotype, suggest that the intron-1 mutation in *VRT-A2b* is the *T. polonicum* species defining polymorphism.

### VRT-A2 expression levels affect spike, glume, grain, and floral organ length in a dosage-dependent manner

We observed that *VRT-A2* was expressed to a higher degree in *P1*^*POL*^ compared to *P1*^*WT*^ NILs in all tested tissues (e.g. developing spikelets, leaves, and anthers). Furthermore, we observed ectopic *VRT-A2* expression in *P1*^*POL*^ tissues that have no detectable expression in wildtype lines (e.g. glumes and grains). These expression profiles were also found in transgenic lines that carry the *VRT-A2b* allele. In these transgenic lines, we found a linear relationship between expression levels and multiple phenotypic traits (R^2^ = 0.76 for spike length; R^2^ = 0.92 for glume length; R^2^ = 0.88 for lemma length; R^2^ = 0.81 for grain length; R^2^ = 0.83 for palea length; all *P* <0.001). The NIL and transgenic data also suggest that outer organs, including glumes and lemmas, are more responsive to changes in *VRT-A2* expression levels than inner organs such as paleae and carpels. Our results suggest that ectopic and higher expression of *VRT-A2* leads to multiple phenotypic effects, including the glume, lemma, and grain length phenotypes, in a dosage dependent manner in polyploid wheat.

In the *P1*^*POL*^ NIL, we detected ectopic expression of *VRT-A2* in tissues that do not show morphological changes (e.g. lemma and grains from floret 4) nor differences in pericarp cell size (e.g. grains from floret 4). Similarly, we only detected an increase in length in paleae from floret 1, but not subsequent florets. This is recapitulated in the transgenic lines with medium copy number of the transgene (Supplementary Table S16). In contrast, the lines with high copy number show significant increases in lemma length (*P* < 0.02) and grain length (*P* ≤ 0.02) within floret 3 and 4, respectively. Similarly, palea length is only significantly different in floret 1 in the *P1*^*POL*^ NIL, whereas paleae are longer up until floret 3 in the high copy number transgenic lines. This suggests that increasing expression levels of *VRT-A2* are necessary to alter organ length in subsequent florets and across different organs. The strongest effects are visible in the basal florets of the spikelets, and only transgenic lines with medium or high copy number exhibit changes in organ length in apical florets, consistent with a basipetal gradient that determines organ length. Both lemma and palea, and the resulting grain, respond to this gradient, although the magnitude of the phenotypic effects (with respect to wildtype or zero copy lines) is stronger in lemmas than in paleae.

A possible explanation for these results is the sequential formation of these organs during spikelet and floral development together with the fact that MADS-box genes act in sequential manner as part of protein complexes (Schwarz-Sommer *et al*., 1992; Goto and Meyerowitz, 1994; Davies *et al*., 1996; Huang *et al*., 1996; Riechmann *et al*., 1996; Egea-Cortines *et al*., 1999; Honma and Goto, 2001). In the wildtype, *VRT2* expression is strongly downregulated during the transition from vegetative meristem to the double ridge stage, presumably to allow floral transition from vegetative to spikelet and floret meristems (Trevaskis *et al*., 2007; Li *et al*., 2019; Li *et al*., 2020). Tetraploid wheat lines with constitutive expression of *VRT2* show significant downregulation of A-, B-, C-, and E-class floral genes at the terminal spikelet stage of spike development (W3.5) (Li *et al*., 2020). In *Arabidopsis*, the SVP-class genes *SVP* and *AGL24* act as repressors of B- and C-class flowering genes (Gregis *et al*., 2013). Thus, normal transition into floral meristems takes place under decreasing SVP levels and increasing levels of A-class (SQUAMOSA) and E-class (SEPALLATA) MADS-box proteins, among others.

Disruption in this balance can lead to increased vegetative characteristics as evidenced in E-class mutants in rice which have leaf-like glumes (rudimentary glumes and sterile lemmas), lemmas, and paleae (Ren *et al*., 2016; Wu *et al*., 2018). Likewise, overexpression of *SVP* genes in wheat (Li *et al*., 2020) and barley (Trevaskis *et al*., 2007) results in a delay or reversion, respectively, of this vegetative to reproductive transition. Li *et al*. (2020) hypothesise that the downregulation of *SVP* genes is necessary given that SVP proteins interfere with SQUAMOSA-SEPALLATA protein complexes that are required for normal spikelet and floral development. This is consistent with the results presented here where higher and ectopic expression of *VRT-A2* in *P1*^*POL*^ and transgenic lines would interfere with the activity of these protein complexes. The magnitude of the response and the tissues affected would be dependent on the level of *SVP* overexpression and the ability of SVP proteins to compete with the sequential MADS-box protein complexes that give rise to the different floral tissue types. We hypothesise that SVP is able to compete more strongly with protein complexes required for glume and lemma development (as shown in Li *et al*. (2020)) and gradually less so with those protein complexes involved in palea development, which include additional MADS-box proteins (e.g. *ALG6-like* genes; (Reinheimer and Kellogg, 2009)). This would explain the dosage-dependent response observed in our study and why we observe the strongest effects in outer and early established organs (e.g. glumes and lemmas) while later developing/differentiating organs (e.g. paleae) are affected only in lines with the highest *VRT-A2* expression.

### Intron 1 motifs are conserved across grasses and may be recognised by repressors

Our results are reminiscent of the pod corn phenotype observed in maize *Tunicate1* (*Tu1*) mutants, in which the grains (kernels) are completely enclosed by elongated glumes. Similar to *P1*, the mutant *Tu1* phenotype is caused by the ectopic expression of the MADS-box gene *ZMM19*, the maize *TaSVP1* homolog and a closely related ortholog of wheat *VRT2*, in the developing maize inflorescence (Supplementary Figure S8; (Han *et al*., 2012; Wingen *et al*., 2012; Schilling *et al*., 2020)). The ectopic expression of *ZMM19*, however, is due to a duplication and rearrangement in the promoter region, whereas our results indicate that the intron-1 sequence plays a key regulatory role in the expression profile of *VRT-A2*.

Numerous MADS-box genes have been shown to contain regulatory sequences within their first introns, including *FLC* in *Arabidopsis* (Sung *et al*., 2006) and *VRN1* in wheat (reviewed in Distelfeld *et al*. (2009)). We thus hypothesise that the 563-bp sequence of the *VRT-A2a* allele, substituted for 160-bp in the *VRT-A2b* allele, contains putative regulatory sequences for establishing the correct expression pattern of the gene. By comparing *VRT2* intron-1 sequences across Poaceae, we identified two distinct motifs (both within the 563-bp region of intron 1) that showed a high degree of sequence conservation across 60 million years of evolution. The absence of the 563-bp intron-1 sequence, as in V*RT-A2b*, results in a misexpression of *VRT-A2*, both in terms of its absolute expression levels and spatiotemporal patterns. It is thus tempting to speculate that either one or both conserved intron-1 motifs allows the binding of proteins or protein complexes that repress *VRT-A2* expression.

Using online databases, we identified significant hits in both intron-1 motifs to members of the LOB-domain family of transcription factors. LOB-domain genes are important for the establishment of boundaries between floral organs and have been shown to be important for glume, lemma, and palea development in several monocot species. In rice, mutations in LOB-domain genes *DEGENERATED HULL 1* (*DH1*; (Li *et al*., 2008)) and *INDETERMINATE GAMETOPHYTE 1* (*OsIG1*; (Zhang *et al*., 2015)) affect glume, lemma, and palea formation, and *OsIG1* affects expression of *SEPALLATA* and *AGL-6* like MADS-box genes. In maize and barley, LOB-domain genes *RAMOSA2 (RA2*; (Bortiri *et al*., 2006)) and *VULGARE ROW-TYPE SPIKE 4 (VRS4*; (Koppolu *et al*., 2013)) restrict inflorescence branching and establish determinacy of spikelet meristems. Strong overexpression of *VRT-A2* in wheat (Li *et al*., 2020), and maize plants with multiple copies of the ectopically expressed *Tu1* allele (Han *et al*., 2012), result in spikelet branching similar to that observed in *ra2* mutants (Bortiri *et al*., 2006). This is consistent with an antagonistic relationship between their activities as first suggested by Han *et al*. (2012). Further investigation, however, will be required to understand if the intron 1 motifs can be recognised by repressors and if LOB-domain proteins play a role in this. We cannot exclude the possibility that the misexpression of *VRT-A2* is caused by the *VRT-A2b* allele. Despite the 160-bp sequence not being homologous to any plant sequence in the NCBI database, it is predicted to contain putative DNA binding motifs for transcription factors (Supplementary Data S1, S2).

The B- and D-homoeologs of *VRT2* also contain the two highly conserved intron-1 motifs, and as such we see no difference in their expression pattern between NILs nor in the transgenic lines. Likewise, no changes in expression of the closest MADS-box ortholog (*TaSVP1*) were detected in *P1* NILs nor *VRT-A2* transgenic lines, similar to the lack of expression differences in closely related MADS-box genes in the maize *Tu1* mutants (Han *et al*., 2012; Wingen *et al*., 2012). This suggests that *VRT-A2* does not regulate its homoeologs or is unable to overcome the presence of the putative repressive protein or protein complex in intron 1 of the B- and D-genome homoeologs. Further work is needed to fully characterise the role of these putative motifs and how they regulate expression of *VRT2*.

### cis-regulatory variation can impact agronomic traits in polyploid wheat

Major loci that control a relatively large proportion of phenotypic variation for quantitative traits have been selected during domestication of diploid plant species (reviewed in Swinnen *et al*. (2016)). Often, the causal variants underlying these phenotypes occur in *cis*-regulatory regions of developmental regulators that affect the level or the spatiotemporal expression profile of transcription factors (Sieburth and Meyerowitz, 1997; Salvi *et al*., 2007; Louwers *et al*., 2009; Studer *et al*., 2011). Selection of *cis*-regulatory variation has also played a pivotal role in shaping polyploid wheat domestication. Examples include the major vernalisation (*VRN1*; (Yan *et al*., 2003)) and photoperiod (*Ppd1*; (Wilhelm *et al*., 2009)) response genes as well as in the major homoeolog pairing *Ph1* locus (Rey *et al*., 2017). All these selected wheat domestication alleles are dominant or semi-dominant, thereby circumventing functional redundancy and allowing the rapid detection of favourable phenotypes.

The *P1*^*POL*^ allele provides a compelling example, where the over- and extended expression of *VRT-A2* results in enhancement of traits of agronomic interest in a dosage-dependent (semi-dominant) manner. This is similar to recent results in maize, where increasing and extending the expression of the MADS-box gene *ZMM28* resulted in improved vegetative and reproductive growth parameters, which impacted positively on yield (Wu *et al*., 2019). Interestingly, the authors discuss how a more subtle over- and extended expression of *ZMM28* using a native maize promoter resulted in more consistent yield benefits and fewer pleiotropic effects compared to promoters with constitutive overexpression. Analogously, overexpression of *VRT-A2* in wheat (Li *et al*., 2020) and related *SVP* genes in barley (Trevaskis *et al*., 2007) and rice (Sentoku *et al*., 2005) using the maize Ubiquitin promoter (in all three studies) resulted in multiple negative pleiotropic effects, including floral reversion. These results highlight how the more subtle changes in expression profiles, through variation in *cis*-regulation, can impact on agronomic traits. Recent work in tomato has shown how a wide range of phenotypic variation for quantitative traits can be engineered by genome editing of transcription factor promoters to generate *cis*-regulatory alleles (Rodriguez-Leal *et al*., 2017). It will be important to determine if engineered *cis*-regulatory variants will overcome functional redundancy and have similar impact on agronomic traits in a polyploid context.

In summary, we identified *VRT2*, a member of the *SVP* family of MADS-box transcription factors, as the gene underlying the *T. polonicum P1* locus in polyploid wheat. An intron-1 sequence rearrangement results in the misexpression of *VRT-A2*, which leads to multiple phenotypic effects in a dosage dependent manner. Allelic variation studies support the intron-1 mutation in *VRT-A2* as the *T. polonicum* species defining polymorphism. The *P1*^*POL*^ allele increases grain weight and other agronomic traits, but not yield, in UK environments. As expression levels of *VRT-A2* are correlated with the magnitude of the phenotypic effects, it is possible that engineering of *VRT2* expression patterns through novel *cis*-regulatory alleles will generate further beneficial quantitative variation for plant breeding.

## Materials and Methods

### Germplasm

To develop *P1* NILs, we crossed *T. polonicum* accession 1100002 to the hexaploid spring wheat cultivar Paragon and the resulting F_1_ was backcrossed four to six times to the Paragon recurrent parent. At each generation, F_1_ lines exhibiting the long-glume phenotype of *T. polonicum* where selected to continue the backcrossing process. After four (BC_4_) or six (BC_6_) backcrosses, BC_n_F_2_ plants were grown and homozygous lines for *P1* selected based on glume length. Bulked seed from the BC_4_F_2_ or BC_6_F_2_ plants were used for subsequent experiments. Accessions of *T. dicoccoides, T. polonicum, T. petropavlovskyi*, and of the *T. aestivum* landrace group ‘Arrancada’ as well as the Watkins collection were obtained from the IPK Genebank, the USDA-ARS National Small Grains Collection (NSGC), the John Innes Centre Germplasm Resources Unit (GRU), the Centre for Genetic Resources (CGN) at Wageningen University, and the International Center for Agricultural Research in the Dry Areas (ICARDA).

### Field experiments and phenotyping

The *P1* NILs were evaluated in six field experiments between 2016 to 2020. Two trials (2016 BC_4_; 2020 BC_4_ and BC_6_) were sown at the John Innes Centre Experimental trials site in Bawburgh, UK (52°37’50.7”N 1°10’39.7”E) and four (2017 BC_4_; 2018, 2019 and 2020 BC_4_ and BC_6_) were sown at The Morley Agricultural Foundation trials site in Morley St Botolph, UK (52°33’15.1”N 1°01’59.2”E). All experiments were sown in autumn (end September-November; except 2020 which was sown in February) as yield-scale plots (6m x 1.2m) in a randomised complete block design (RCBD) with five replications and sown by grain number for comparable plant densities aiming for 275 seeds*m^−2^. Developmental traits were evaluated throughout the growing period and a 10-ear sample was collected at harvest for the assessment of spike, floret and grain characteristics (marked ‘10ES’ in Supplementary Table S1). Spike length was measured as the distance between the peduncle-rachis junction and tip of the terminal floret. Plot yield, hectolitre weight, and grain moisture were measured during harvest on board the combine (Zürn 150). Final grain yield was determined per plot after adjustment to 15% grain moisture. Grain morphometric measurements were analysed using the MARVIN grain analyser (GTA Sensorik GmbH, Neubrandenburg, Germany) using *∼* 400 grains of the combined grain samples.

### Spike dissection and organ measurements (NILs and transgenic lines)

We measured organ size of the *P1*^*WT*^ and *P1*^*POL*^ NILs by sampling three spikes from five field blocks per NIL grown in 2019 at Morley. The spikes were dissected by hand and all organs (glume, lemma, palea, and grain) were placed on PCR film (Cat No.: AB0580, Thermofisher) from bottom (position 0 in Fig. 1E-F) to top (position 20 in Fig 1E-F) of the spike. The PCR films with the organs were scanned using a standard Ricoh photocopier (settings: greyscale, 600dpi). The resulting images were analysed using the Fiji “analyse particles” function, restricting analysis to particles of 0.1-5 cm^2^ area (Schindelin *et al*., 2012). Fiji measures particles from top-left to bottom-right of the image, thus allowing us to match position of the organ along the spike with the Fiji measurements retrospectively. To measure organ size in the transgenic T_0_ lines (grown in 1 L pots under 16 hours light at 20°C and 8 hours darkness at 15°C in a controlled environment room), we hand dissected organs from two main spikes per plant. The organs were measured and analysed as described for the NILs above.

### 3D scanning of spikes and morphometric grain extraction

Fifteen mature spikes from both *P1* NILs grown in 2019 at Morley were used for μCT scanning (three spikes from five field blocks per NIL). Scanning conditions were as described in Hughes *et al*. (2019). Feature extraction from the scans was performed using previously developed MATLAB-based software (Hughes *et al*., 2017) using the following setup parameters (SE=7, voxel size=68.8, minSize=10,000 and watershed=false). The features extracted were length (calculated using the major axis of the whole grain), width, and depth (the major and minor axis of a cross-section respectively, found by selecting the grain’s midpoint), volume (a complete connected pixel count per grain), and grain counts for each spike. More than 750 grains were measured per genotype. The data were checked for false positives by first removing outliers that were identified using the 0.025 upper and lower percentiles of the data. Additionally, for added robustness, manual checks were performed.

### Grain developmental timecourse

The *P1* NILs grown in 2018 and 2019 at Morley were used for the grain developmental time courses. For this, we tagged 70 ears per NIL over five replicated blocks in the field at ear emergence (spike fully emerged and peduncle just visible), as described in Brinton *et al*. (2017). Ten spikes per NIL were collected at five (2018) and six (2019) different timepoints. These were ear emergence, 3, 9, 16, and 22 days post anthesis (dpa) in 2018, while in 2019, the timepoints included ear emergence, anthesis (here measured as anther extrusion), 7, 14, 21, and 28 dpa. Grain measurements were performed as described in Brinton *et al*. (2017).

### Cell size measurements

We measured cell size of mature grains from *P1*^*WT*^ and *P1*^*POL*^ NILs collected from three field blocks grown at Morley in 2019. Within each block, we sampled three spikes and from each spike we sampled grains from florets 2 and 4 of the two central spikelets. In total, this resulted in 18 grains per genotype per floret position (2 grains x 3 spikes x 3 field blocks). Dry grain samples were mounted crease-down onto 12.5 mm diameter aluminium pin stubs using double-sided 12 mm adhesive carbon discs (Agar Scientific Ltd, Stansted, Essex). The stubs were then sputter coated with approximately 15nm gold in a high-resolution sputter coater (Agar Scientific Ltd) and transferred to a Zeiss Supra 55 VP FEG scanning electron microscope (Zeiss SMT, Germany). The samples were viewed at 3kV with a magnification of 1500x and digital TIFF files were stored. The surface of each grain was imaged in the top, middle and bottom thirds of the grain (excluding the embryo; Supplementary Figure S6) with three images taken in each section (nine images total per grain). Cell length was measured manually using the Fiji distribution of ImageJ (Schindelin *et al*., 2012). For statistical analyses, only images with ≥30 cell measurements were used. For each image, the median cell length was calculated. The image medians were then used to calculate a median cell length value for each section (bottom/middle/top) of each grain.

### Genetic Mapping of P1

For fine-mapping, we generated a set of BC_4_ and BC_6_ recombinant inbred lines (RILs) derived from the *P1* NILs. In the first round we identified 17 BC_4_F_2_ heterozygous recombinant lines between markers *S1* and *S9*. We screened twelve BC_4_F_3_ progeny for each line to identify homozygous recombinants, which were phenotyped for glume length (Supplementary Table S7). To further define the *P1* interval, we screened an additional 1867 BC_6_F_2_ plants heterozygous across the *S2* and *S7* interval. We identified 64 independent homozygous recombinants between markers *S2* and *S10*, which were phenotyped for glume length and genotyped with a further 21 markers (Supplementary Table S8). The eight critical recombinants (Supplementary Table S9) were grown at the John Innes Centre Experimental trials site and phenotyped for height, grain weight, spike length and grain morphometrics.

To test the isogenic nature of the BC_4_ *P1* NILs, we used the Axiom 35k Breeders’ Array (Allen *et al*., 2017). The array showed that 98.7% of markers (32839) were monomorphic between the NILs, with 418 polymorphisms between the NILs. More than 65% of the polymorphisms (272) were located on chromosome 7A, while the remaining were distributed evenly across other chromosomes. To generate markers, we performed exome-capture of an accession of *T. polonicum* (idPlant: 27422, GRU Store Code: T1100002), wildtype Paragon, and wildtype Langdon. These three samples were exome-sequenced in a pool of eight samples on a single Illumina HiSeq2000 lane following published protocols (Krasileva *et al*., 2017). This generated 27919048, 30795964, and 30683631 reads for the three lines, respectively. The reads were mapped to the RefSeqv1.0 (International Wheat Genome Sequencing *et al*., 2018) assembly using bwa-0.7.15 (bwa mem -t 8 -M; (Li and Durbin, 2009; Li, 2013)). The resulting SAM file was converted to BAM format using samtools-1.3.1 (samtools view -b - h; (Li *et al*., 2009)) and sorted by chromosome position (samtools sort). Optical and PCR duplicates were marked using picard-1.134 (picard MarkDuplicates MAX_FILE_HANDLES_FOR_READ_ENDS_MAP=1024 VALIDATION_STRINGENCY=LENIENT http://broadinstitute.github.io/picard/). Single nucleotide polymorphisms (SNPs) were called for chromosome 7A with freebayes-1.1.0 (freebayes -0 -t; (Garrison and Marth, 2012)), and filtered using bcftools-1.3.1 (bcftools filter; (Li and Durbin, 2009)). Lastly, the vcf file was compressed with bgzip, indexed with tabix-0.2.6 (tabix -p vcf; (Li and Durbin, 2009)) before extracting relevant data in a user-friendly format with bcftools-1.3.1 (bcftools query -H -f ‘%CHROM\t%POS\t%REF\t%ALT{0}\t%QUAL\t%INFO/DP\t%INFO/RO\t%INFO/AO{0}[\t%GT\t %DP\t%RO\t%AO{0}]\n’). The SNPs were filtered for polymorphisms between *T. polonicum* and the two cultivars Paragon and Langdon. These putative SNPs were used to design KASP markers using PolyMarker (Ramirez-Gonzalez *et al*., 2015). KASP assays were validated in the parental NILs and then used for genetic mapping of *P1* as indicated in Supplementary Tables S7-S8.

#### PCR markers

The mapping populations were genotyped as described in Trick *et al*. (2012), with the following changes: 2 µl DNA (10-40 ng) was mixed with 2 µl of mastermix (2 µl PACE (Standard ROX; 3CR Bioscience) with 0.056 µl primer assay) for a total reaction volume of 4 µl. All PACE markers used for map-based cloning are listed in Supplementary Table S18. Standard PCR as well as qRT-PCR primers, their annealing temperatures and amplicon sizes are listed in Supplementary Table S19.

#### TraesCS7A02G175200 gene model

Using the expVIP browser (Borrill *et al*., 2016; Ramirez-Gonzalez *et al*., 2018), expression of *TraesCS7A02G175200* showed high expression in young seedlings (vegetative plants with 1 cm long spikes). The corresponding transcriptome data (Zadoks growth stage 30; (Choulet *et al*., 2014)) was downloaded, and aligned to the genomic RefSeqv1.0 assembly using HiSat2 v2.1.0 (hisat2 -p 16; (Kim *et al*., 2015)). The SAM file was converted to BAM format, sorted, and optical duplicates were marked as described for the exome capture data above. The depth of reads was measured using samtools-1.3.1 (samtools depth -a). The RNA-seq data supports the *TraesCS7A02G175200*.*1* gene model and its predicted untranslated regions (UTRs; Supplementary Figure S7B).

#### Phylogenetic analysis of StMADS11-like family

Amino acid sequences of StMADS11 and related proteins were aligned in MEGA X using MUSCLE with default settings (Gap open penalty: -2.9; Gap extension penalty: 0; Hydrophobicity multiplier: 1.2; Clustering method: UPGMA; (Edgar, 2004b, a; Kumar *et al*., 2018; Schilling *et al*., 2020)). The alignment was then used to create a neighbor-joining tree in MEGA X (Test of phylogeny: 1000 bootstraps; Model: Poisson; Rates among sites: Uniform; Gaps/Missing Data: Pairwise deletion; (Zuckerkandl and Pauling, 1965; Felsenstein, 1985; Saitou and Nei, 1987; Kumar *et al*., 2018)). A list of all proteins used for the alignment can be found in Supplementary Table S20. The alignment and tree files are deposited at Dryad (https://datadryad.org/stash/share/a6HM2SGbQyigaK7r2AnYc3TFao1kF9kN8C1QzScBsUU).

### Haplotype variation of TraesCS7A02G175200

We sequenced the promoter (2299 bp), *TraesCS7A02G175200* genomic sequence (5591 bp, exon and introns) and 1857 bp downstream of the termination codon (9747 bp) in the *P1*^*POL*^ NIL using primers detailed in Supplementary Table S19. We also sequenced the 5591 bp exon-intron sequences of *TraesCS7A02G175200* in six *T. polonicum*, two *T. petropavlovskyi*, and four ‘Arrancada’ accessions (Supplementary Table S12) using primers listed in Supplementary Table S19.

We developed a PCR marker (primers S37_Fwd and S37_Rev; Supplementary Table S19) to determine the presence of either the 563-bp (*VRT-A2a* allele) or the 160-bp (*VRT-A2b* allele) intron-1 rearrangement in a large diversity panel. We used this marker to assay the intron-1 status of 70 wild emmer (*T. dicoccoides*), 103 hexaploid landraces, 4 durum, 23 *T. polonicum*, 2 *T. petropavlovskyi* and 7 ‘Arrancada’ landrace accessions (Supplementary Table S11, S12). We also used available genome sequences of 16 hexaploid (Walkowiak *et al*., 2020) and 3 tetraploid (Avni *et al*., 2017; Maccaferri *et al*., 2019; Walkowiak *et al*., 2020) cultivars and accessions to characterise *TraesCS7A02G175200* across its promoter, exon-intron sequences and 3’ untranslated region (Supplementary Table S10). We also evaluated the wider haplotype of the *P1* NILs, 16 hexaploid and 1 tetraploid cultivar, 7 *T. polonicum*, 2 *T. petropavlovskyi* and 7 ‘Arrancada’ landrace accessions using 14 markers spanning the 7A physical region (Supplementary Table S13). Details of the accessions used are listed in Supplementary Tables S11 and S12.

### Phylogenetic footprinting

We extracted intron-1 sequences of *VRT2* orthologs from barley (HORVU7Hr1G036130), *Brachypodium* (Bradi1g45812), rice (Os06g0217300), maize (GRMZM5G814279) and sorghum (SORBI_3010G085400) and used these, alongside *TraesCS7A02G175200*, as query sequences in the mVISTA program (http://genome.lbl.gov/vista/index.shtml).

### Intron-1 motif discovery

The phylogenetic footprinting analysis revealed two conserved sequence peaks between wheat, barley, *Brachypodium*, rice, maize, and sorghum. These sequences, plus some flanking sequence (71 and 84 bp for both regions, respectively), were aligned using T-Coffee with default settings (https://www.ebi.ac.uk/Tools/msa/tcoffee/; (Notredame *et al*., 2000; Madeira *et al*., 2019)). We defined the motifs using the following approach: a nucleotide was considered conserved if it was identical in 5 out of the 6 species (83%). A maximum of four nucleotides with lower conservation was tolerated, provided the neighbouring sequences were again highly conserved (83%). This yielded a 34 and 69 bp sequence, which were designated Motif 1 and Motif 2, respectively (Supplementary Data Set S1). When tolerating only a single nucleotide with low conservation (<83%), a highly conserved 16 and 20 bp sequence was detected within Motif 1 and Motif 2, respectively (Supplementary Data Set S1). In addition, within Motif 2, we detected a 6 bp highly conserved palindromic sequence that we designated “palindrome” (Supplementary Data Set S1). As there are no homologous sequences to the *VRT-A2b* allele 160-bp rearrangement, we divided the sequence into three “motifs”, constituting the two sections made of matching flanking DNA and the leftover sequence (Supplementary Data Set S2).

### Transcription factor binding site analysis of predicted motifs

To search for possible transcription factor binding sites within the predicted motif sequences we used three different tools. The sequences were queried one by one using the Binding Site Prediction tool from PlantRegMap (http://plantregmap.gao-lab.org/binding_site_prediction.php) with default settings for the *Arabidopsis thaliana, Oryza sativa*, and *Zea mays* databases respectively (Tian *et al*., 2020). Next, we queried the *Arabidopsis thaliana, Oryza sativa*, and *Zea mays* databases of PlantPan3.0 using the TF/TFBS Search tool with a q-value cut-off of 0.05. Lastly, we used the Tomtom tool of MEME Suite 5.2.0 (http://meme-suite.org/tools/tomtom) to query the *Arabidopsis thaliana* DAP motifs database from O’Malley *et al*. (2016) as well as the ‘JASPAR CORE (2018) plants’ database (Khan *et al*., 2018) with default settings.

### RNA extraction

BC_4_ NILs were grown in 11 cm^2^ pots (1 L volume) in ‘John Innes Cereal Mix’ (40% Medium Grade Peat, 40% Sterilised Soil, 20% Horticultural Grit, 1.3 kg*m^-3^ PG Mix 14-16-18 + Te Base Fertiliser, 1 kg*m^-3^ Osmocote Mini 16-8-11 2 mg + Te 0.02% B, Wetting Agent, 3 kg*m^-3^ Maglime, 300 g*m^-3^ Exemptor) under long day conditions (16 h light : 8 h dark) in the glasshouse. Tissues were harvested, immediately placed into 2 ml tubes in liquid nitrogen and stored at -80°C until needed. For meristem tissues, samples were dissected using a stereo microscope (Leica MZ16) and processed as above. Details of tissues sampled are presented in Supplementary Table S14. For transgenic plants, we sampled flag leaves, glumes, and grains at 21 days post anthesis.

The grain samples were homogenized using mortar and pestle with liquid nitrogen. All other tissues were homogenized in a SPEX CertiPrep 2010-230 Geno/Grinder (Cat No.: 12605297, Fischer Scientific) using 5 mm steel beads (Cat No.: 69989, Qiagen); tubes were shaken in 20 sec bursts at 1500 rpm, then immediately transferred back into liquid nitrogen. Depending on the tissue type, this was repeated up to two times.

RNA was extracted using three different methods depending on the tissue:

(a) For young spikes (up until Floret primordium stage W3.5), we used the Qiagen RNeasy Plant Mini Kit (Cat No.: 74904, Qiagen) with RLT buffer according to the manufacturer’s protocol, as it enables recovery of RNA from small input samples. DNA digestion was performed using the RNase-Free DNase Set (Cat No.: 79254, Qiagen) according to the manufacturer’s protocol.
(b) For all other non-grain tissues, we used the Spectrum Plant Total RNA kit (Cat No.: STRN250-1KT, Sigma), following Protocol A of the manufacturer’s protocol and using 750 μL of Binding Solution. DNA digestion was performed using the On-Column DNase I Digestion Set (Cat No.: DNASE70-1SET, Sigma) according to the manufacturer’s protocol.
(c) For grain samples, 500 μL of RNA extraction buffer (0.1 M Tris pH 8.0, 5 mM EDTA pH 8.0, 0.1 M NaCl, 0.5% SDS; autoclaved) with 1% β-Mercaptoethanol (Cat No.: M3148, Merck) and 100 μL of Ambion Plant RNA Isolation Aid (Cat No.: AM9690, Thermofisher) were added to each sample, before vortexing. Tissue debris as well as polysaccharides and polyphenols were pelleted at 13000 rpm for 10 min in a microcentrifuge. The supernatant was transferred to a new 1.5 mL tube, before adding 500 μL of Acid Phenol:Chloroform:IAA (125:24:1) (Cat No.: AM9720, Thermofisher). The tubes were shaken in a SPEX CertiPrep 2010-230 Geno/Grinder for 10 min at 500 rpm, then placed in a microcentrifuge at 13000 rpm for 15 min to separate the organic and aqueous components. The supernatant (aqueous phase) was transferred to a new 1.5 mL tube with 500 μL of Chloroform (Cat No.: C/4960/PB17, FisherScientific). The tubes were inverted 10 times and then placed in a microcentrifuge for 15 min at 13000 rpm. The supernatant was transferred to a new 1.5 mL tube with 360 μL of Isopropanol (Cat No.: P/7500/PC17, FisherScientific) and 45 μL 3 M Sodium Acetate (pH 5.2). The tube was inverted 10 times to mix the solution, before placing at 4°C for 1 hour to precipitate RNA. The RNA was pelleted in a microcentrifuge at 4°C by spinning for 30 min at 13000 rpm. The supernatant was carefully tipped off to not lose the pellet. The tubes were then washed twice with 70% Ethanol (Cat No.: 20821.330, VWR) and centrifuged between washes at 13000 rpm for 5 min at 4°C. The supernatant was then carefully discarded and remaining droplets of Ethanol removed using a pipette tip, before adding 100 μL of nuclease-free water (Cat No.: AM9937, Thermofisher).

### Quantitative real-time reverse-transcription PCR (qRT-PCR)

RNA was reverse transcribed using M-MLV reverse transcriptase (Cat No.: 28025013, Thermofisher) according to the manufacturer’s protocol. For the qRT-PCR reactions, LightCycler 480 SYBR Green I Master Mix (Roche Applied Science, UK) was used according to the manufacturer’s protocol. The reactions were run in a LightCycler 480 instrument (Roche Applied Science, UK) under the following conditions: 5 min at 95 °C; 45 cycles of 10 s at 95 °C, 15 s at 62 °C, 30 s at 72 °C; dissociation curve from 60 °C to 95 °C to determine primer specificity. All reactions were performed with three technical replicates per sample and using *TaActin* as reference gene (Li *et al*., 2019). Relative gene expression was calculated using the 2^-ΔΔCt^ method (Livak and Schmittgen, 2001) with a common calibrator so that values are comparable across genes, tissues and developmental stages. All primers used in qRT-PCR are listed in Supplementary Table S19.

### Construct assembly

A modified version of the GoldenGate (MoClo) compatible level 2 vector pGoldenGreenGate-M (pGGG-M) as described in Hayta *et al*. (2019) was used in this study. The pGGG-AH-L2P2 acceptor plasmid is comprised of the hygromycin resistance gene (hpt) containing the Cat1 intron driven by the rice actin1 (*OsAct1*) promoter for *in planta* selection and a LacZ-MCS flanked by two *Bsa*I sites at MoClo position 2 with standardised overhangs to accept basic (level 0) components. In brief, the *T. polonicum VRT-A2* promoter (2299 bp), genomic sequence (5585 bp), 1000 bp downstream of STOP codon, and NOS terminator (8916 bp total) were cloned into pGGG-AH-L2P2 using standard Golden Gate MoClo assembly (Werner *et al*., 2012), resulting in construct pGGG-AH-VRT-A2 (Supplementary Figure S10). Several *Bsa*I and *Bbs*I sites had to be domesticated to make the *T. polonicum VRT-A2* sequence suitable for Golden Gate MoClo assembly, including 3 sites in the promoter (C4174T, G4549A, C5755T), 1 site in exon 1 (C6126T; V47V), 1 site in an intronic MITE (C8377T), and 1 site in exon 3 (T9477C; L106L). Six nucleotides from a partial LINE in intron 5 were omitted by mistake from the genomic sequence (hence 5585 bp from start to termination codon instead of 5591 bp). The construct was electroporated into the hypervirulent *Agrobacterium tumefaciens* strain AGL1 (Lazo *et al*., 1991) containing the helper plasmid pAL155 (additional *Vir*G gene). Standard inoculums of *Agrobacterium* (Tingay *et al*., 1997) were prepared as described in Hayta *et al*. (2019).

### Wheat transformation

Hexaploid wheat c.v. ‘Fielder’ was transformed using the previously described method by Hayta *et al*. (2019). In brief, under aseptic conditions wheat immature embryos were isolated, pre-treated by centrifugation, inoculated with *A. tumefaciens* AGL1 containing pGGG-AH-VRT-A2 and co-cultivated for 3 days. Wheat callus proliferation, shoot regeneration, and rooting were carried out under a stringent hygromycin selection regime before the regenerated plantlets were transferred from *in vitro* to soil and acclimatised. Transgenesis was confirmed by *hpt* gene PCR; transgene copy number analysis was performed using Taqman qPCR and probe (Hayta *et al*., 2019). The values obtained were used to calculate copy number according to published methods (Livak and Schmittgen, 2001). Based on this copy number determination we defined T_0_ lines as zero, low (1 to 2 copies of pGGG-AH-VRT-A2), medium (4-5 copies of pGGG-AH-VRT-A2) and high (9 or more copies of pGGG-AH-VRT-A2) copy number lines (Supplementary Table S15).

### Statistical Analyses

*Field experiments*: To determine the differences between the *P1*^*POL*^ and *P1*^*WT*^ NILs, we performed ANOVA on the multiple field phenotypic data in RStudio (v1.3.1056). For the overall analysis we included block (nested in location), genotype, location, and the genotype*location interaction in the model. For the analysis of individual locations, we used a simple two-way ANOVA including block and genotype. For the BC_6_ RILs, we determined the effect of the *VRT-A2* allele on height, TGW, spike length, grain width and length using a two-way ANOVA using block and the *VRT-A2* genotype in the model. For glume length, RILs were assigned as having a normal or long-glume phenotype using a *post hoc* Dunnett’s test to compare with the *P1*^*POL*^ and *P1*^*WT*^ controls.

#### Spike dissection

We used glume measurements from spikelets 1 (basal) to 20 (apical) to determine the differences between the *P1*^*POL*^ and *P1*^*WT*^ NILs. We did not include datapoints from spikelets 21 to 23 as these were only found in very few instances (balanced between genotype) and made up less than 2.5% of the total grain data (50 datapoints excluded from 2117 total grain values). Likewise, we did not include 80 datapoints from florets 5, which between them represented 3.8% of grain data and were found only in spikelets 4 to 13 in both NILs. Given that the experimental unit is the field plot to which the genotype was randomised within each block, we analysed the data using a split-plot ANOVA in which the ‘Spikelet’ was nested within the ‘Genotype*Block’ interaction. The ANOVA therefore included the following terms: Block, Genotype, Spikelet and Genotype*Spikelet interactions, with *F* statistic and *P* values calculated based on the ‘Block*Genotype’ error term (for Block and Genotype) or the Residual error term for the other factors.

We analysed the grain, lemma, and palea data from spikelets 1 to 20 and florets 1 to 4 to determine the differences between the *P1* NILs. Given that the florets are nested in the spikelet, and the spikelet is nested within the genotype*block interaction (i.e. the florets and spikelets are not randomly assigned), we analysed the data as a split-split plot design using the corresponding error terms for calculating the *F* statistic and *P* values. This model included the block, genotype, spikelet, floret, and corresponding interaction terms. We also performed individual ANOVAs for each floret position using the same model as above, with the exception that we excluded floret as a factor.

#### Grain development timecourse

Each block at every timepoint consisted of *c*. 100 grains (10 spikes x 10 grains) per NIL. The grain morphometrics were averaged across the 100 grains (as they were considered to be subsamples) to yield a single value per timepoint, resulting in 5 datapoints per NIL per timepoint. We performed a two-way ANOVA with Block and Genotype in the model to determine whether *P1* affects grain morphometrics at the sampled timepoints.

#### Cell size measurements

We analysed the data independently for floret 2 and 4 using a three-way ANOVA including Block, Genotype, Block*Genotype, Section, and the Genotype*Section interaction. Given the significant Genotype*Section interactions, we explored differences between *P1* genotypes for each section using Tukey multiple comparison as implemented in RStudio (v1.3.1056).

#### Expression

We evaluated differences in expression levels of *VRT2* and *MADS22* homoeologs by performing t-tests between the 2^-ΔΔCt^ expression values of *P1*^*POL*^ and *P1*^*WT*^ NILs for each individual tissue*timepoint comparison.

#### Phenotypes in transgenic lines

To evaluate differences in phenotype between the four categories of transgenic lines (zero, low, medium, and high copy number lines; Supplementary Table S16) we performed one-way ANOVAs for each floret position including ‘transgene copy number’ as the single factor. Given that ‘transgene copy number’ was significant for all phenotypes (glume, lemma, palea, and grain length) and across all florets, we performed Tukey multiple comparison tests to determine differences between the four ‘transgene copy number’ categories as well as Dunnett tests against the zero copy number control lines (Supplementary Table S16).

#### Correlation of phenotype and expression in transgenic T0 lines

We calculated the Pearson’s correlation coefficient between *VRT-A2* expression (in flag leaf, glume, and grain) and phenotypic traits (internode length, spike length, glume length, lemma length, grain length, palea length) in R (Supplementary Table S17). We used geom_smooth(method = “lm”) to plot the regression line and 95% confidence interval.

### Accession numbers

Sequence data from this article can be found in the EMBL/GenBank data libraries under accession numbers MW289820 -MW289831 (*TraesCS7A02G175200* sequence for *T. polonicum, T. petropavlovskyi*, and Arrancada accessions), and MW307955 – MW307976 for *T. dicoccoides* and Watkins accessions (Supplementary Table S11, S12). Construct pGGG-AH-VRT-A2 has been deposited in Addgene (ID 163703) and the sequence is available (NCBI accession MW289819). Seed of the BC_6_ *P1*^*POL*^ NIL is available from the JIC GRU (Code WM0013). Full-size images of spikes and spikelets of the *T. polonicum, T. petropavlovskyi*, and ‘Arrancada’ accessions used in this study and in Figure 3B-C are deposited at Dryad (https://datadryad.org/stash/share/a6HM2SGbQyigaK7r2AnYc3TFao1kF9kN8C1QzScBsUU).

### Supplemental Data files

**Supplementary Data Set 1**: Analysis of *VRT-A2* intron-1 conserved sequences across grass species.

**Supplementary Data Set 2**: Analysis of putative transcription factor binding sites within *VRT-A2* intron 1

## Supporting information

Supplementary Tables and Data Sets

## Acknowledgements

This work was supported by the UK Biotechnology and Biological Sciences Research Council (BBSRC) through the grant BB/S016945/1, BB/S016538/1, the Designing Future Wheat (BB/P016855/1), the National Capability in Plant Phenotyping (BBS/E/W/0012844A), Genes in the Environment (BB/P013511/1) Institute Strategic Programmes and Core Capability Grant. We are grateful to Dr. Harold Bockelman (NSGC), Dr. Ulrike Lohwasser (IPK Genebank), Dr. Noam Chayut (GRU), Dr. Noortje Bas (CGN), and Dr. Athanasios Tsivelikas (ICARDA) for providing germplasm. We thank the JIC Bioimaging facility and staff for their contribution to this publication. We also thank Phil Robinson from the Scientific Photography at JIC for taking high quality images of spikes and spikelets. We thank the JIC Field Trials and Horticultural Services teams for technical support in field and glasshouse experiments and Karen Askew for uCT support.

## Author Contributions

JS, NMA, and CU conceived the study. JS generated the NILs and mapping populations together with PS. NMA performed most experiments. AP and OH genotyped Watkins, *T. dicoccoides*, and *T. polonicum* accessions using a PCR marker and sequenced amplicons. YC helped with the analysis of *VRT-A2* alleles and haplotypes. AEB dissected spikelets of NILs, phenotyped and assessed the data. The grains of these lines were imaged by JFB and EB using scanning electron microscopy, with JFB analysing the data. MS designed and created the pGGG-AH-VRT-A2 construct; SH transformed cv. Fielder with the construct, cultivated the plants and performed copy number analysis on them. JS performed field experiments and phenotyping. TF and PC helped with plant husbandry, phenotyping, and data collection. MC, JD, and CN performed CT-scans of field-grown spikelets and analysed the data. JFB, NMA and CU created the figures. CU performed statistical analysis of all data. NMA and CU wrote the manuscript. All authors have read and approved the manuscript.

## References

Akond, A.S.M.G., and Watanabe, N. (2005). Genetic variation among portuguese landraces of ‘Arrancada’ wheat and Triticum petropavlovskyi by AFLP-based assessment. Genet Resour Crop Ev 52, 619–628.

Akond, A.S.M.G.M., Watanabe, N., and Furuta, Y. (2008). Comparative genetic diversity of Triticum aestivum - Triticum polonicum introgression lines with long glume and Triticum petropavlovskyi by AFLP-based assessment. Genet Resour Crop Ev 55, 133–141.

Allen, A.M., Winfield, M.O., Burridge, A.J., Downie, R.C., Benbow, H.R., Barker, G.L., Wilkinson, P.A., Coghill, J., Waterfall, C., Davassi, A., et al. (2017). Characterization of a Wheat Breeders’ Array suitable for high-throughput SNP genotyping of global accessions of hexaploid bread wheat (Triticum aestivum). Plant Biotechnol J 15, 390–401.

Avni, R., Nave, M., Barad, O., Baruch, K., Twardziok, S.O., Gundlach, H., Hale, I., Mascher, M., Spannagl, M., Wiebe, K., et al. (2017). Wild emmer genome architecture and diversity elucidate wheat evolution and domestication. Science 357, 93–97.

Biffen, R.H. (2009). Mendel’s laws of inheritance and wheat breeding. The Journal of Agricultural Science 1, 4–48.

Borrill, P., Ramirez-Gonzalez, R., and Uauy, C. (2016). expVIP: a customizable RNA-seq data analysis and visualization platform. Plant Physiol 170, 2172–2186.

Bortiri, E., Chuck, G., Vollbrecht, E., Rocheford, T., Martienssen, R., and Hake, S. (2006). ramosa2 encodes a LATERAL ORGAN BOUNDARY domain protein that determines the fate of stem cells in branch meristems of maize. Plant Cell 18, 574–585.

Brinton, J., Simmonds, J., Minter, F., Leverington-Waite, M., Snape, J., and Uauy, C. (2017). Increased pericarp cell length underlies a major quantitative trait locus for grain weight in hexaploid wheat. New Phytol 215, 1026–1038.

Charles, M., Tang, H., Belcram, H., Paterson, A., Gornicki, P., and Chalhoub, B. (2009). Sixty million years in evolution of soft grain trait in grasses: emergence of the softness locus in the common ancestor of Pooideae and Ehrhartoideae, after their divergence from Panicoideae. Mol Biol Evol 26, 1651–1661.

Chen, P.D., D.J., L., G.Z., P., L.L., Q., and L., H. (1988). The chromosome constitution of three endemic hexaploid wheats in Western China. In 7th Int. Wheat Genet. Symp, M. T.E. and K. R.M.D., eds (Cambridge, UK), pp. 75–80.

Chen, Q., Y., S., and Y., D. (1985). Cytogenetic studies on interspecific hybrids of Xinjiang wheat. Acta Agron Sin 11, 23–28.

Choulet, F., Alberti, A., Theil, S., Glover, N., Barbe, V., Daron, J., Pingault, L., Sourdille, P., Couloux, A., Paux, E., et al. (2014). Structural and functional partitioning of bread wheat chromosome 3B. Science 345, 1249721.

Davies, B., Egea-Cortines, M., de Andrade Silva, E., Saedler, H., and Sommer, H. (1996). Multiple interactions amongst floral homeotic MADS box proteins. EMBO J 15, 4330–4343.

Deriano, L., and Roth, D.B. (2013). Modernizing the Nonhomologous End-Joining repertoire: alternative and classical NHEJ share the stage. Annual Review of Genetics 47, 433–455.

Distelfeld, A., Li, C., and Dubcovsky, J. (2009). Regulation of flowering in temperate cereals. Curr Opin Plant Biol 12, 178–184.

Edgar, R.C. (2004a). MUSCLE: multiple sequence alignment with high accuracy and high throughput. Nucleic Acids Res 32, 1792–1797.

Edgar, R.C. (2004b). MUSCLE: a multiple sequence alignment method with reduced time and space complexity. BMC Bioinformatics 5, 113.

Egea-Cortines, M., Saedler, H., and Sommer, H. (1999). Ternary complex formation between the MADS-box proteins SQUAMOSA, DEFICIENS and GLOBOSA is involved in the control of floral architecture in Antirrhinum majus. EMBO J 18, 5370–5379.

Engledow, F.L. (1920). The inheritance of glume-length and grain-length in a wheat cross. Journal of Genetics 10, 109–134.

Felsenstein, J. (1985). Confidence limits on phylogenies: an approach using the bootstrap. Evolution 39, 783–791.

Frazer, K.A., Pachter, L., Poliakov, A., Rubin, E.M., and Dubchak, I. (2004). VISTA: computational tools for comparative genomics. Nucleic Acids Res 32, W273–279.

Garrison, E., and Marth, G. (2012). Haplotype-based variant detection from short-read sequencing. ArXiv 1207.3907.

Goto, K., and Meyerowitz, E.M. (1994). Function and regulation of the Arabidopsis floral homeotic gene PISTILLATA. Genes Dev 8, 1548–1560.

Gregis, V., Andres, F., Sessa, A., Guerra, R.F., Simonini, S., Mateos, J.L., Torti, S., Zambelli, F., Prazzoli, G.M., Bjerkan, K.N., et al. (2013). Identification of pathways directly regulated by SHORT VEGETATIVE PHASE during vegetative and reproductive development in Arabidopsis. Genome Biol 14, R56.

Han, J.J., Jackson, D., and Martienssen, R. (2012). Pod corn is caused by rearrangement at the Tunicate1 locus. Plant Cell 24, 2733–2744.

Hayta, S., Smedley, M.A., Demir, S.U., Blundell, R., Hinchliffe, A., Atkinson, N., and Harwood, W.A. (2019). An efficient and reproducible Agrobacterium-mediated transformation method for hexaploid wheat (Triticum aestivum L.). Plant Methods 15, 121.

Honma, T., and Goto, K. (2001). Complexes of MADS-box proteins are sufficient to convert leaves into floral organs. Nature 409, 525–529.

Huang, H., Tudor, M., Su, T., Zhang, Y., Hu, Y., and Ma, H. (1996). DNA binding properties of two Arabidopsis MADS domain proteins: binding consensus and dimer formation. Plant Cell 8, 81–94.

Hughes, N., Oliveira, H.R., Fradgley, N., Corke, F.M.K., Cockram, J., Doonan, J.H., and Nibau, C. (2019). µCT trait analysis reveals morphometric differences between domesticated temperate small grain cereals and their wild relatives. Plant J 99, 98–111.

Hughes, N., Askew, K., Scotson, C.P., Williams, K., Sauze, C., Corke, F., Doonan, J.H., and Nibau, C. (2017). Non-destructive, high-content analysis of wheat grain traits using X-ray micro computed tomography. Plant Methods 13, 76.

International Wheat Genome Sequencing, C., Appels, R., Eversole, K., Feuillet, C., Keller, B., Rogers, J., Stein, N., Pozniak, C.J., Stein, N., Choulet, F., et al. (2018). Shifting the limits in wheat research and breeding using a fully annotated reference genome. Science 361, eaar7191.

Kane, N.A., Agharbaoui, Z., Diallo, A.O., Adam, H., Tominaga, Y., Ouellet, F., and Sarhan, F. (2007). TaVRT2 represses transcription of the wheat vernalization gene TaVRN1. Plant J 51, 670–680.

Kane, N.A., Danyluk, J., Tardif, G., Ouellet, F., Laliberte, J.F., Limin, A.E., Fowler, D.B., and Sarhan, F. (2005). TaVRT-2, a member of the StMADS-11 clade of flowering repressors, is regulated by vernalization and photoperiod in wheat. Plant Physiol 138, 2354–2363.

Kent, T., Mateos-Gomez, P.A., Sfeir, A., and Pomerantz, R.T. (2016). Polymerase theta is a robust terminal transferase that oscillates between three different mechanisms during end-joining. Elife 5, e13740.

Khan, A., Fornes, O., Stigliani, A., Gheorghe, M., Castro-Mondragon, J.A., van der Lee, R., Bessy, A., Cheneby, J., Kulkarni, S.R., Tan, G., et al. (2018). JASPAR 2018: update of the open-access database of transcription factor binding profiles and its web framework. Nucleic Acids Res 46, D260–D266.

Kim, D., Langmead, B., and Salzberg, S.L. (2015). HISAT: a fast spliced aligner with low memory requirements. Nat Methods 12, 357–360.

Koppolu, R., Anwar, N., Sakuma, S., Tagiri, A., Lundqvist, U., Pourkheirandish, M., Rutten, T., Seiler, C., Himmelbach, A., Ariyadasa, R., et al. (2013). Six-rowed spike4 (Vrs4) controls spikelet determinacy and row-type in barley. Proc Natl Acad Sci U S A 110, 13198–13203.

Kosuge, K., Watanabe, N., and Kuboyama, T. (2010). Recombination around the P locus for long glume phenotype in experimental introgression lines of Triticum aestivum -Triticum polonicum. Genet Resour Crop Ev 57, 611–618.

Krasileva, K.V., Vasquez-Gross, H.A., Howell, T., Bailey, P., Paraiso, F., Clissold, L., Simmonds, J., Ramirez-Gonzalez, R.H., Wang, X., Borrill, P., et al. (2017). Uncovering hidden variation in polyploid wheat. Proc Natl Acad Sci U S A 114, E913–E921.

Kumar, S., Stecher, G., Li, M., Knyaz, C., and Tamura, K. (2018). MEGA X: Molecular Evolutionary Genetics Analysis across computing platforms. Mol Biol Evol 35, 1547–1549.

Langdale, J.A., Irish, E.E., and Nelson, T.M. (1994). Action of the Tunicate locus on maize floral development. Developmental Genetics 15, 176–187.

Lazo, G.R., Stein, P.A., and Ludwig, R.A. (1991). A DNA transformation-competent Arabidopsis genomic library in Agrobacterium. Biotechnology (N Y) 9, 963–967.

Li, A., Zhang, Y., Wu, X., Tang, W., Wu, R., Dai, Z., Liu, G., Zhang, H., Wu, C., Chen, G., et al. (2008). DH1, a LOB domain-like protein required for glume formation in rice. Plant Mol Biol 66, 491–502.

Li, C., Lin, H., Chen, A., Lau, M., Jernstedt, J., and Dubcovsky, J. (2019). Wheat VRN1, FUL2 and FUL3 play critical and redundant roles in spikelet development and spike determinacy. Development 146, dev175398.

Li, H. (2013). Aligning sequence reads, clone sequences and assembly contigs with BWA-MEM. ArXiv 1303.3997.

Li, H., and Durbin, R. (2009). Fast and accurate short read alignment with Burrows-Wheeler transform. Bioinformatics 25, 1754–1760.

Li, H., Handsaker, B., Wysoker, A., Fennell, T., Ruan, J., Homer, N., Marth, G., Abecasis, G., Durbin, R., and Genome Project Data Processing, S. (2009). The Sequence Alignment/Map format and SAMtools. Bioinformatics 25, 2078–2079.

Li, K., Debernardi, J.M., Li, C., Lin, H., Zhang, C., and Dubcovsky, J. (2020). Interactions between SQUAMOSA and SVP MADS-box proteins regulate meristem transitions during wheat spike development. bioRxiv, https://doi.org/10.1101/2020.1112.1101.405779.

Livak, K.J., and Schmittgen, T.D. (2001). Analysis of relative gene expression data using real-time quantitative PCR and the 2^−ΔΔCt^ method. Methods 25, 402–408.

Louwers, M., Bader, R., Haring, M., van Driel, R., de Laat, W., and Stam, M. (2009). Tissue- and expression level-specific chromatin looping at maize b1 epialleles. Plant Cell 21, 832–842.

Maccaferri, M., Harris, N.S., Twardziok, S.O., Pasam, R.K., Gundlach, H., Spannagl, M., Ormanbekova, D., Lux, T., Prade, V.M., Milner, S.G., et al. (2019). Durum wheat genome highlights past domestication signatures and future improvement targets. Nat Genet 51, 885–895.

Madeira, F., Madhusoodanan, N., Lee, J., Tivey, A.R.N., and Lopez, R. (2019). Using EMBL-EBI Services via web interface and programmatically via web services. Curr Protoc Bioinformatics 66, e74.

Mangelsdorf, P.C., and Galinat, W.C. (1964). The Tunicate locus in maize dissected and reconstituted. Proc Natl Acad Sci U S A 51, 147–150.

Matsumura, S. (1950). Linkage study in wheat. II. P-linkage group and the manifold effects of the P gene. Jpn J Genet 25, 111–118.

Mayor, C., Brudno, M., Schwartz, J.R., Poliakov, A., Rubin, E.M., Frazer, K.A., Pachter, L.S., and Dubchak, I. (2000). VISTA : visualizing global DNA sequence alignments of arbitrary length. Bioinformatics 16, 1046–1047.

Millet, E. (1986). Relationships between grain weight and the size of floret cavity in the wheat spike. Annals of Botany 58, 417–423.

Notredame, C., Higgins, D.G., and Heringa, J. (2000). T-Coffee: a novel method for fast and accurate multiple sequence alignment. J Mol Biol 302, 205–217.

O’Malley, R.C., Huang, S.C., Song, L., Lewsey, M.G., Bartlett, A., Nery, J.R., Galli, M., Gallavotti, A., and Ecker, J.R. (2016). Cistrome and epicistrome features shape the regulatory DNA landscape. Cell 165, 1280–1292.

Okamoto, Y., and Takumi, S. (2013). Pleiotropic effects of the elongated glume gene P1 on grain and spikelet shape-related traits in tetraploid wheat. Euphytica 194, 207–218.

Percival, J. (1921). The wheat plant: a monograph. (London: Duckworth and Co.).

Ramirez-Gonzalez, R.H., Uauy, C., and Caccamo, M. (2015). PolyMarker: A fast polyploid primer design pipeline. Bioinformatics 31, 2038–2039.

Ramirez-Gonzalez, R.H., Borrill, P., Lang, D., Harrington, S.A., Brinton, J., Venturini, L., Davey, M., Jacobs, J., van Ex, F., Pasha, A., et al. (2018). The transcriptional landscape of polyploid wheat. Science 361, eaar6089.

Reineke, A.R., Bornberg-Bauer, E., and Gu, J. (2011). Evolutionary divergence and limits of conserved non-coding sequence detection in plant genomes. Nucleic Acids Res 39, 6029–6043.

Reinheimer, R., and Kellogg, E.A. (2009). Evolution of AGL6-like MADS box genes in grasses (Poaceae): ovule expression is ancient and palea expression is new. Plant Cell 21, 2591–2605.

Ren, D., Rao, Y., Leng, Y., Li, Z., Xu, Q., Wu, L., Qiu, Z., Xue, D., Zeng, D., Hu, J., et al. (2016). Regulatory role of OsMADS34 in the determination of glumes fate, grain yield, and quality in rice. Front Plant Sci 7, 1853.

Rey, M.D., Martin, A.C., Higgins, J., Swarbreck, D., Uauy, C., Shaw, P., and Moore, G. (2017). Exploiting the ZIP4 homologue within the wheat Ph1 locus has identified two lines exhibiting homoeologous crossover in wheat-wild relative hybrids. Mol Breed 37, 95.

Riechmann, J.L., Wang, M., and Meyerowitz, E.M. (1996). DNA-binding properties of Arabidopsis MADS domain homeotic proteins APETALA1, APETALA3, PISTILLATA and AGAMOUS. Nucleic Acids Res 24, 3134–3141.

Rodgers, K., and McVey, M. (2016). Error-prone repair of DNA double-strand breaks. J Cell Physiol 231, 15–24.

Rodriguez-Leal, D., Lemmon, Z.H., Man, J., Bartlett, M.E., and Lippman, Z.B. (2017). Engineering quantitative trait variation for crop improvement by genome editing. Cell 171, 470–480 e478.

Saitou, N., and Nei, M. (1987). The neighbor-joining method: a new method for reconstructing phylogenetic trees. Mol Biol Evol 4, 406–425.

Salvi, S., Sponza, G., Morgante, M., Tomes, D., Niu, X., Fengler, K.A., Meeley, R., Ananiev, E.V., Svitashev, S., Bruggemann, E., et al. (2007). Conserved noncoding genomic sequences associated with a flowering-time quantitative trait locus in maize. Proc Natl Acad Sci U S A 104, 11376–11381.

Schilling, S., Kennedy, A., Pan, S., Jermiin, L.S., and Melzer, R. (2020). Genome-wide analysis of MIKC-type MADS-box genes in wheat: pervasive duplications, functional conservation and putative neofunctionalization. New Phytol 225, 511–529.

Schimmel, J., van Schendel, R., den Dunnen, J.T., and Tijsterman, M. (2019). Templated insertions: a smoking gun for polymerase theta-mediated end joining. Trends Genet 35, 632–644.

Schindelin, J., Arganda-Carreras, I., Frise, E., Kaynig, V., Longair, M., Pietzsch, T., Preibisch, S., Rueden, C., Saalfeld, S., Schmid, B., et al. (2012). Fiji: an open-source platform for biological-image analysis. Nat Methods 9, 676–682.

Schwarz-Sommer, Z., Hue, I., Huijser, P., Flor, P.J., Hansen, R., Tetens, F., Lonnig, W.E., Saedler, H., and Sommer, H. (1992). Characterization of the Antirrhinum floral homeotic MADS-box gene deficiens: evidence for DNA binding and autoregulation of its persistent expression throughout flower development. EMBO J 11, 251–263.

Sentoku, N., Kato, H., Kitano, H., and Imai, R. (2005). OsMADS22, an STMADS11-like MADS-box gene of rice, is expressed in non-vegetative tissues and its ectopic expression induces spikelet meristem indeterminacy. Mol Genet Genomics 273, 1–9.

Sieburth, L.E., and Meyerowitz, E.M. (1997). Molecular dissection of the AGAMOUS control region shows that cis elements for spatial regulation are located intragenically. Plant Cell 9, 355–365.

Studer, A., Zhao, Q., Ross-Ibarra, J., and Doebley, J. (2011). Identification of a functional transposon insertion in the maize domestication gene tb1. Nat Genet 43, 1160–1163.

Sung, S., He, Y., Eshoo, T.W., Tamada, Y., Johnson, L., Nakahigashi, K., Goto, K., Jacobsen, S.E., and Amasino, R.M. (2006). Epigenetic maintenance of the vernalized state in Arabidopsis thaliana requires LIKE HETEROCHROMATIN PROTEIN 1. Nat Genet 38, 706–710.

Swinnen, G., Goossens, A., and Pauwels, L. (2016). Lessons from domestication: targeting cis-regulatory elements for crop improvement. Trends Plant Sci 21, 506–515.

Tian, F., Yang, D.C., Meng, Y.Q., Jin, J., and Gao, G. (2020). PlantRegMap: charting functional regulatory maps in plants. Nucleic Acids Res 48, D1104–D1113.

Tingay, S., McElroy, D., Kalla, R., Fieg, S., Wang, M.B., Thornton, S., and Brettell, R. (1997). Agrobacterium tumefaciens-mediated barley transformation. Plant Journal 11, 1369–1376.

Trevaskis, B., Tadege, M., Hemming, M.N., Peacock, W.J., Dennis, E.S., and Sheldon, C. (2007). Short vegetative phase-like MADS-box genes inhibit floral meristem identity in barley. Plant Physiol 143, 225–235.

Trick, M., Adamski, N.M., Mugford, S.G., Jiang, C.C., Febrer, M., and Uauy, C. (2012). Combining SNP discovery from next-generation sequencing data with bulked segregant analysis (BSA) to fine-map genes in polyploid wheat. BMC Plant Biol 12, 14.

van Kregten, M., de Pater, S., Romeijn, R., van Schendel, R., Hooykaas, P.J., and Tijsterman, M. (2016). T-DNA integration in plants results from polymerase-theta-mediated DNA repair. Nat Plants 2, 16164.

Waddington, S.R., Cartwright, P.M., and Wall, P.C. (1983). A quantitative scale of spike initial and pistil development in barley and wheat. Annals of Botany 51, 119–130.

Walkowiak, S., Gao, L., Monat, C., Haberer, G., Kassa, M.T., Brinton, J., Ramirez-Gonzalez, R.H., Kolodziej, M.C., Delorean, E., Thambugala, D., et al. (2020). Multiple wheat genomes reveal global variation in modern breeding. Nature, https://doi.org/10.1038/s41586-41020-42961-x.

Watanabe, N., and Imamura, I. (2002). Genetic control of long glume phenotype in tetraploid wheat derived from Triticum petropavlovskyi Udacz. et Migusch. Euphytica 128, 211–217.

Watanabe, N., Yotani, Y., and Furuta, Y. (1996). The inheritance and chromosomal location of a gene for long glume in durum wheat. Euphytica 91, 235–239.

Watanabe, N., Bannikova, S.V., and Goncharov, N.P. (2004). Inheritance and chromosomal location of the gene for long glume phenotype found in Portuguese landraces of hexaploid wheat, “Arrancada”. J. Genet. & Breed. 58, 273–278.

Werner, S., Engler, C., Weber, E., Gruetzner, R., and Marillonnet, S. (2012). Fast track assembly of multigene constructs using Golden Gate cloning and the MoClo system. Bioeng Bugs 3, 38–43.

Wilhelm, E.P., Turner, A.S., and Laurie, D.A. (2009). Photoperiod insensitive Ppd-A1a mutations in tetraploid wheat (Triticum durum Desf.). Theor Appl Genet 118, 285–294.

Wingen, L.U., Munster, T., Faigl, W., Deleu, W., Sommer, H., Saedler, H., and Theissen, G. (2012). Molecular genetic basis of pod corn (Tunicate maize). Proc Natl Acad Sci U S A 109, 7115–7120.

Wu, D., Liang, W., Zhu, W., Chen, M., Ferrandiz, C., Burton, R.A., Dreni, L., and Zhang, D. (2018). Loss of LOFSEP transcription factor function converts spikelet to leaf-like structures in rice. Plant Physiol 176, 1646–1664.

Wu, J., Lawit, S.J., Weers, B., Sun, J., Mongar, N., Van Hemert, J., Melo, R., Meng, X., Rupe, M., Clapp, J., et al. (2019). Overexpression of zmm28 increases maize grain yield in the field. Proc Natl Acad Sci U S A 116, 23850–23858.

Yan, L., Loukoianov, A., Tranquilli, G., Helguera, M., Fahima, T., and Dubcovsky, J. (2003). Positional cloning of the wheat vernalization gene VRN1. Proc Natl Acad Sci U S A 100, 6263–6268.

Zhang, J., Tang, W., Huang, Y., Niu, X., Zhao, Y., Han, Y., and Liu, Y. (2015). Down-regulation of a LBD-like gene, OsIG1, leads to occurrence of unusual double ovules and developmental abnormalities of various floral organs and megagametophyte in rice. J Exp Bot 66, 99–112.

Zuckerkandl, E., and Pauling, L. (1965). Evolutionary divergence and convergence in proteins. In Evolving Genes and Proteins, V. Bryson and H.J. Vogel, eds (Academic Press), pp. 97–166.

