## Supplementary Files for "Increased and ectopic expression of *Triticum polonicum VRT-A2* underlies elongated glumes and grains in hexaploid wheat in a dosage-dependent manner"

### Supplementary Figures

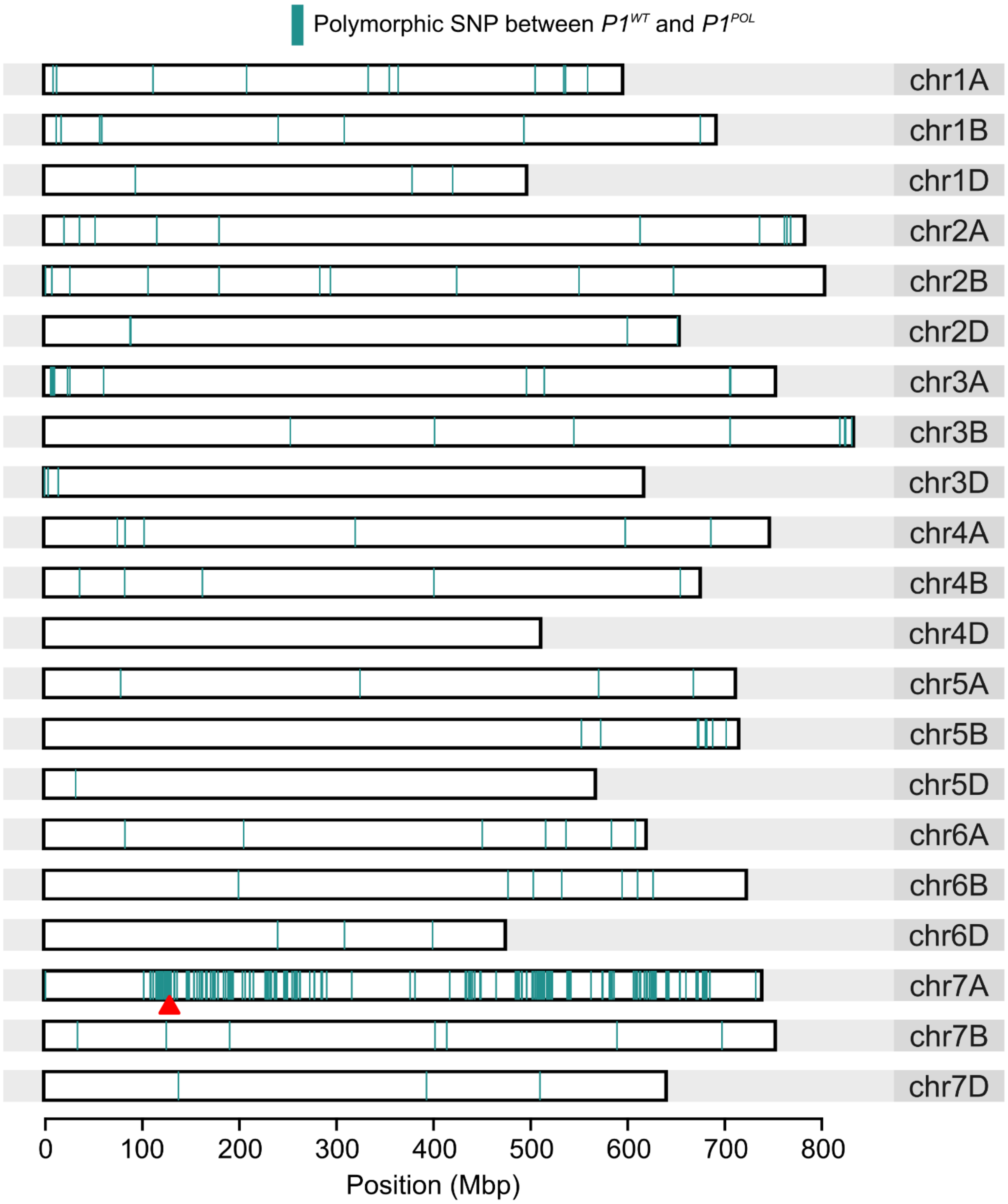


**Supplementary Figure S1** (Supports Figure 1, 2): Genotyping of $BC_4$ *P1* NILs using Breeders' 35K Axiom Array. Polymorphic markers between $P1^{WT}$ and $P1^{POL}$ are indicated as green lines. Physical positions of markers were determined using the IWGSC RefSeqv1.0 assembly. The position of *VRT-A2* is marked by a red triangle.

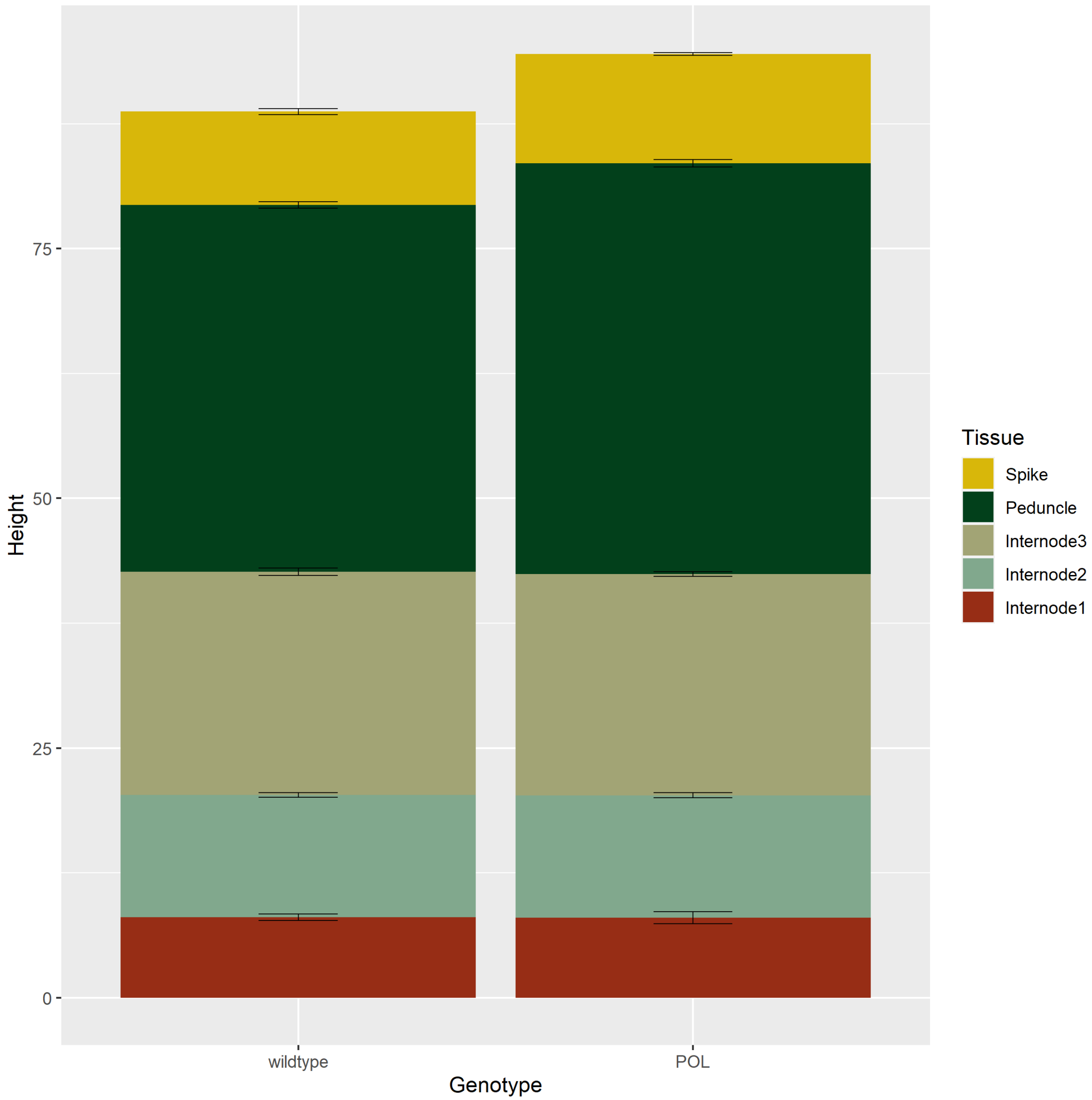


**Supplementary Figure S2** (Supports Table 1)**:** Length of internodes, peduncles, and spikes in $P1^{WT}$ (N=7) and $P1^{POL}$ (N=5) NILs grown in the field in 2017. The three internodes are not significantly different between the NILs, whereas the peduncle is significantly different ($P$ < 0.001) and the spike, in this experiment, is borderline ($P$ <0.06) non-significant. Error bars represent mean ± SEM.

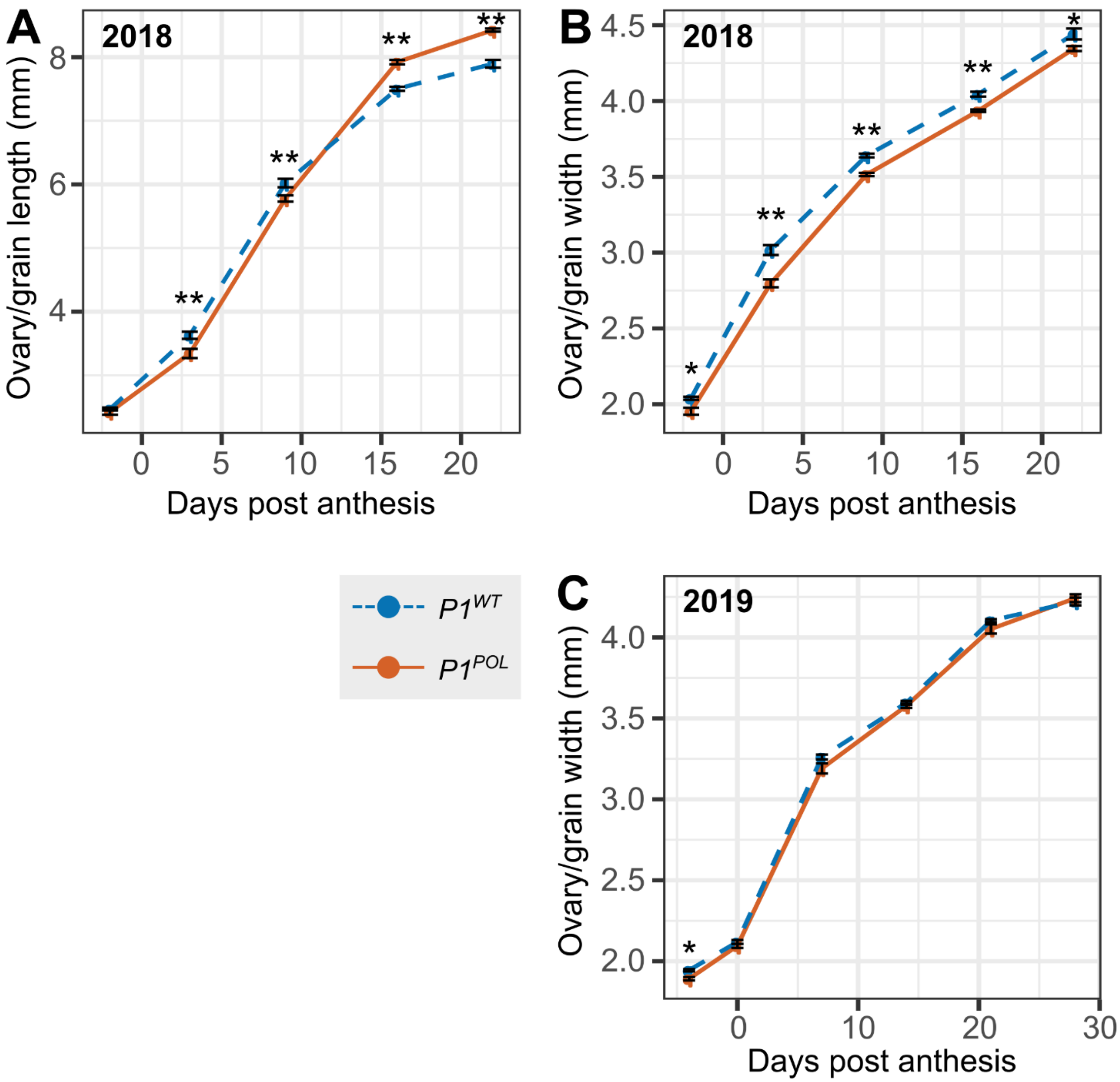


**Supplementary Figure S3** (Supports Figure 1): Timecourse tracking ovary/grain length and width of field-grown *P1*$^{WT}$ and *P1*$^{POL}$ NILs from 2018 (**A, B**) and 2019 (**C**). Length from 2019 is presented in Figure 1D (N=50 per timepoint per genotype). Error bars represent mean ± SEM. *, $P < 0.05$; **, $P < 0.01$; ***, $P < 0.001$.

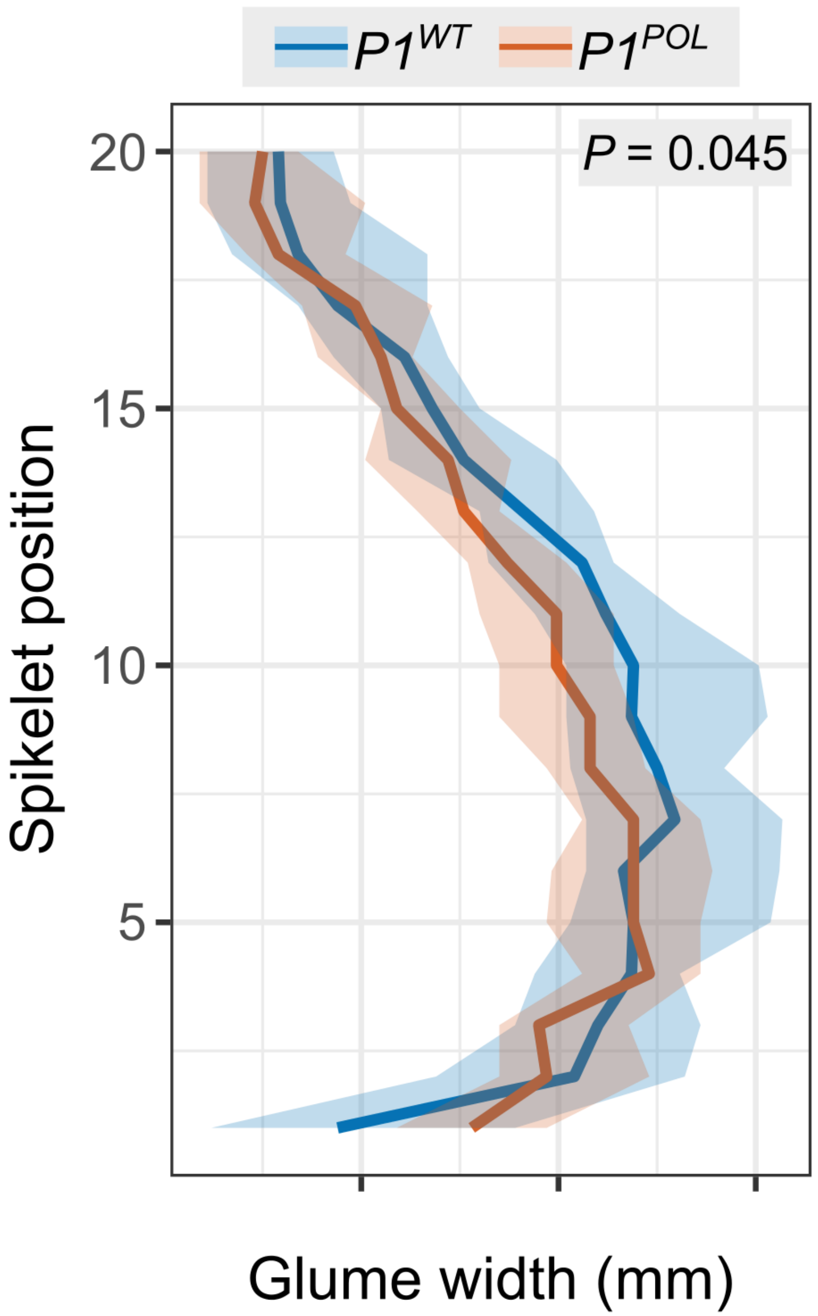


**Supplementary Figure S4** (Supports Figure 1): Glume width along spikes of $P1^{WT}$ and $P1^{POL}$ NILs. Positions are numbered from basal to apical spikelets. Bold line represents the median value, ribbon represents the interquartile range (N=15 spikes).

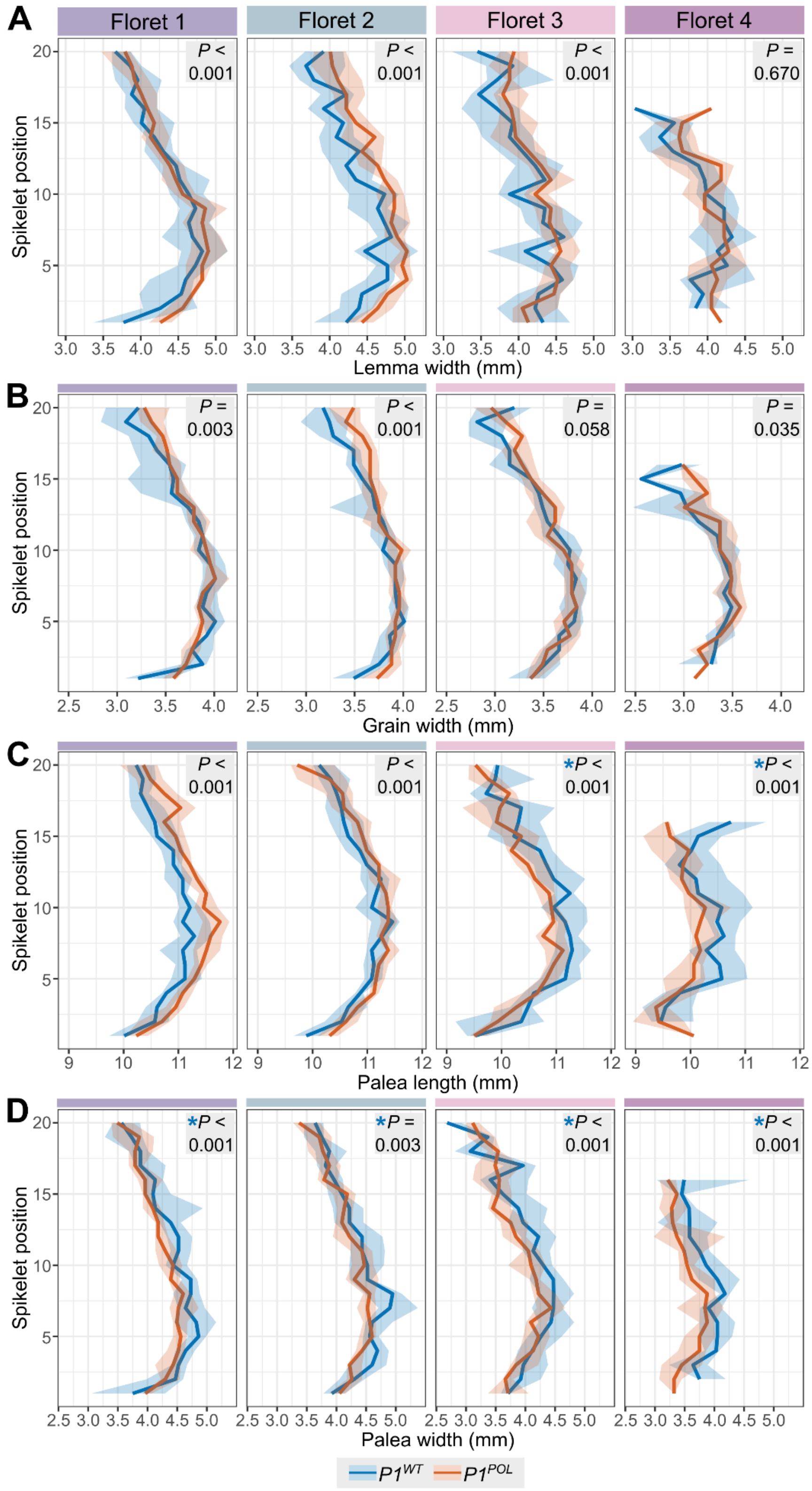


**Supplementary Figure S5** (Supports Figure 1)**:** Lemma width (**A**), grain width (**B**), palea length (**C**), and palea width (**D**) at each floret position along $P1^{WT}$ and $P1^{POL}$ NILs spikes. Spikelet positions are numbered from basal to apical spikelets. Bold line represents the median value, ribbon represents the interquartile range (N=15 spikes).

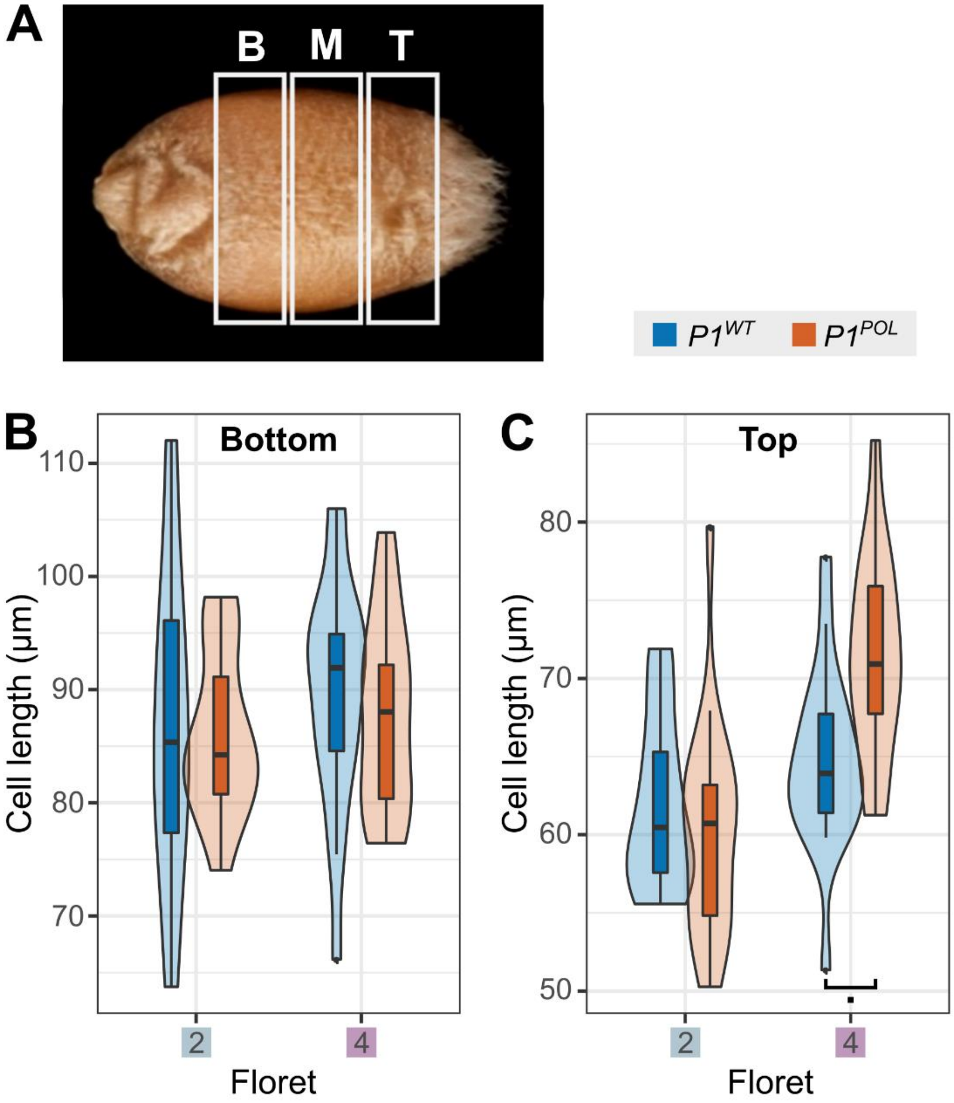


**Supplementary Figure S6** (Supports Figure 1): Pericarp cell length in *P1* NILs. (**A**) Illustration of grain sections used for SEM, including bottom (B), middle (M) and top (T) sections. Pericarp cell length from bottom (**B**) and top (**C**) sections of grains from floret 2 and floret 4 for the $P1^{WT}$ and $P1^{POL}$ NILs (n= 18 grains). In (B) and (C), the box represents the middle 50% of data with the borders of the box representing the 25th and 75th percentile. The horizonal line in the middle of the box represents the median. Whiskers represent the minimum and maximum values, unless a point exceeds 1.5 times the inter-quartile range in which case the whisker represents this value and values beyond this are plotted as single points (outliers). *, $P < 0.05$.

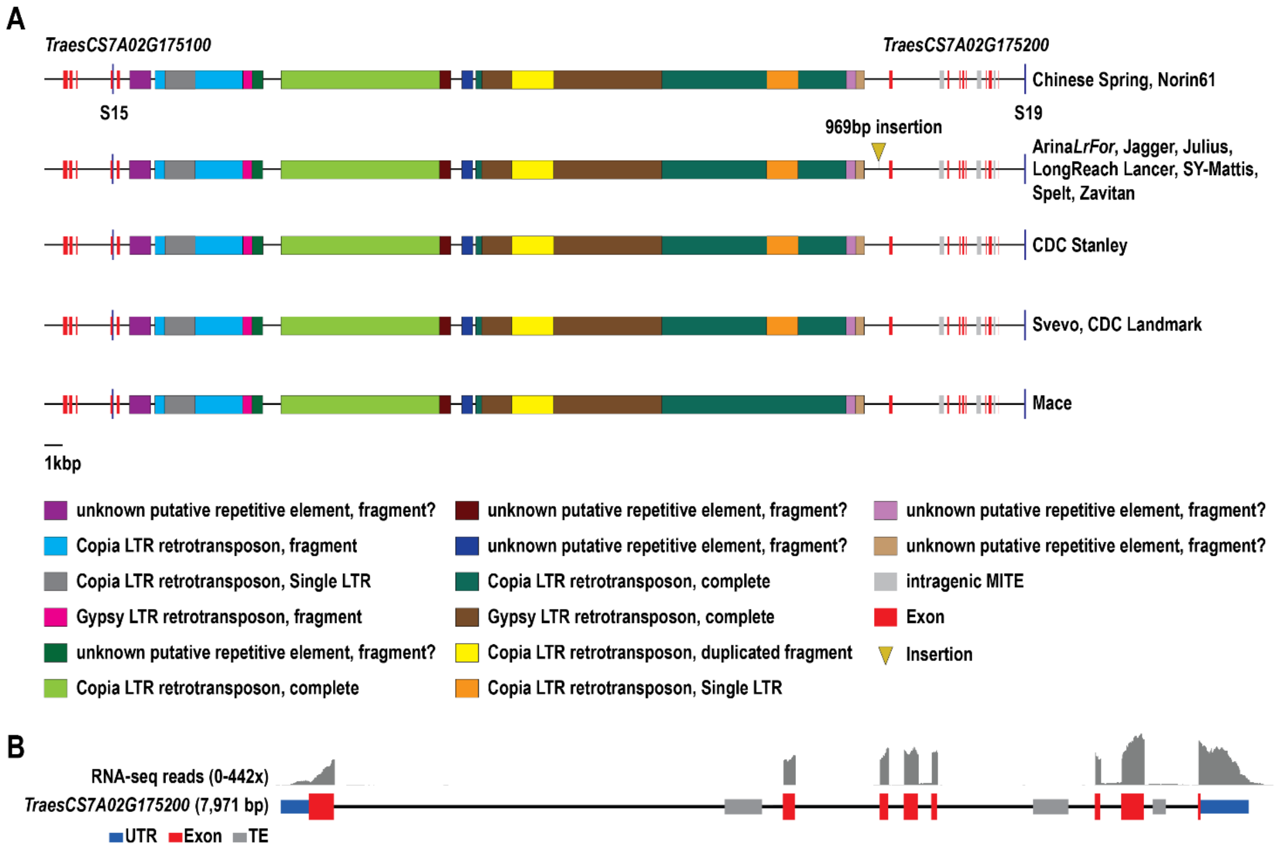


**Supplementary Figure S7** (Supports Figure 2, 3): Characterisation of *P1* critical interval. (**A**) Annotation of physical interval between markers *S15* and *S19* in pangenome cultivars (Walkowiak *et al.*, 2020). (**B**) RNA-Seq coverage of *TraesCS7A02G175200* across predicted exon-intron structure.

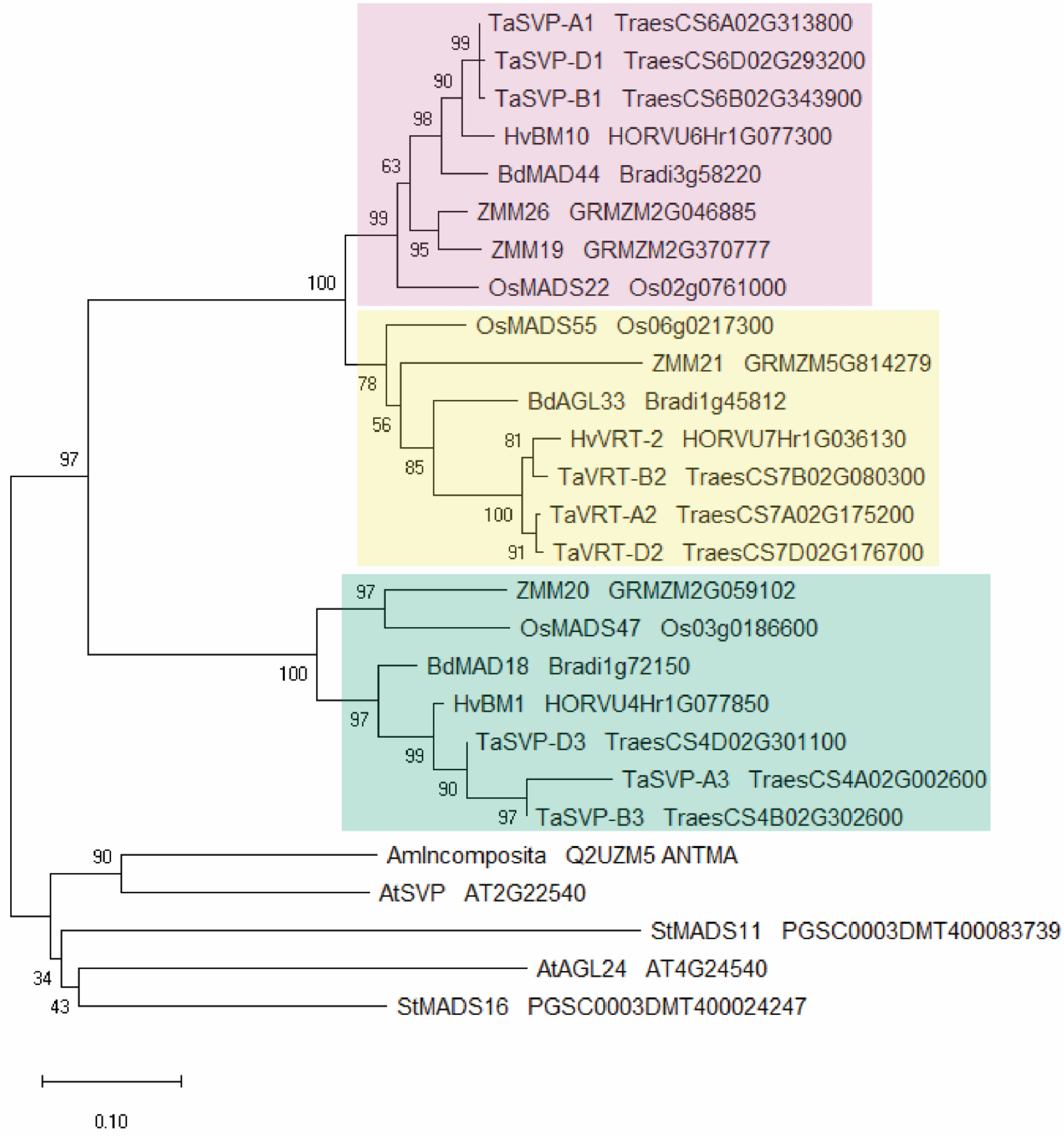


**Supplementary Figure S8** (Supports Figure 4): Phylogenetic tree of StMADS11-like proteins from dicots and monocots made using neighbor-joining method and rooted at the midpoint (Saitou and Nei, 1987). Three SVP-like genes are present in grasses (four in maize), with VRT2 (yellow box) and TaSVP1 (pink box) being more similar to each other than to TaSVP3 (cyan box; (Schilling *et al.*, 2020)). Numbers next to branches indicate the percentage of replicate trees in which the associated proteins clustered together in the bootstrap test (1000 replicates; (Felsenstein, 1985)). Branch lengths were calculated using the Poisson correction method in units of amino acid substitutions per site (Zuckerkandl and Pauling, 1965).

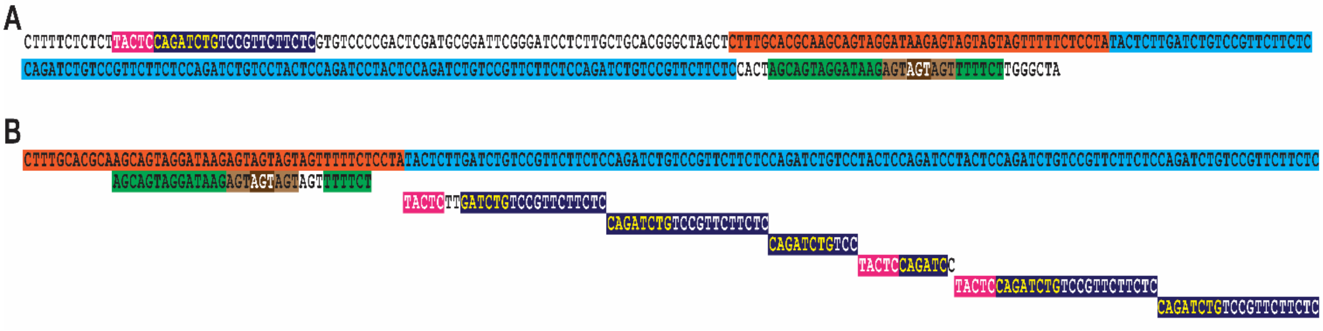


**Supplementary Figure S9** (Supports Figure 2, 3): Examination of the 160-bp rearrangement within the *VRT-A2b* allele. (**A**) The 160-bp sequence rearrangement (highlighted in orange and blue) is shown with 5’ and 3’ flanking sequence. The rearrangement has no overall homology to other plant sequences. However, two sequences flanking the rearrangement (highlighted pink/blue and green/brown) occur within the 160-bp rearrangement. (**B**) The 160-bp rearrangement is shown on a single line. Matching flanking sequences are aligned underneath, with nucleotide mismatches displayed in black font with no highlight. The 160-bp sequence can be divided into two sections. The first section (orange) contains a sequence found 3’ of the 160-bp rearrangement (green; AGT units in brown) with a single polymorphism: the 3’ flanking sequence contains three direct repeats of an AGT triplet, while the sequence within the first section has four repeats of this AGT triplet. The second section (blue) consists entirely of tandem copies (full and partial) of a sequence occurring 5’ of the re-arrangement (pink/blue; yellow font represents a palindromic sequence). The entire sequence (25 bp) appears twice within the second section, albeit one copy contains a CA/TT polymorphism. An additional four incomplete copies occur within the second section. These matching sequences account for 145 out of the 160 nucleotides within the *VRT-A2b* rearrangement.

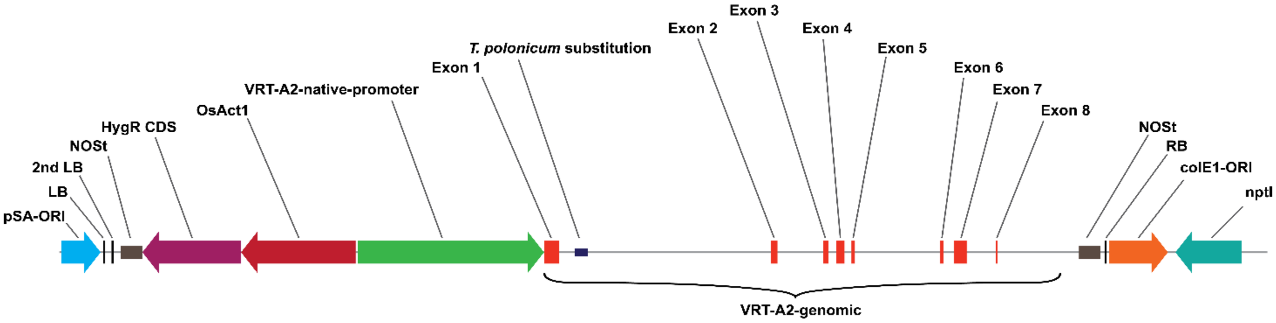


**Supplementary Figure S10** (Supports Figure 5): *VRT-A2* complementation construct. The promoter and genomic sequence of *T. polonicum VRT-A2* were cloned into the pGGG-M vector, in opposite orientation to the Hygromycin resistance coding sequence (HygR CDS) driven by the rice *Actin* promoter for selection (Addgene #163703).
